# Measuring Stimulus Information Transfer Between Neural Populations through the Communication Subspace

**DOI:** 10.1101/2024.11.06.622283

**Authors:** Oren Weiss, Ruben Coen-Cagli

**Affiliations:** Department of Systems and Computational Biology, Albert Einstein College of Medicine, Bronx, NY 10461, USA; Dominick P. Purpura Department of Neuroscience, Albert Einstein College of Medicine, Bronx, NY 10461, USA; Department of Ophthalmology and Visual Sciences, Albert Einstein College of Medicine, Bronx, NY 10461, USA

## Abstract

Sensory processing arises from the communication between neural populations across multiple brain areas. While the widespread presence of neural response variability shared throughout a neural population limits the amount of stimulus-related information those populations can accurately represent, how this variability affects the interareal communication of sensory information is unknown. We propose a mathematical framework to understand the impact of neural population response variability on sensory information transmission. We combine linear Fisher information, a metric connecting stimulus representation and variability, with the framework of communication subspaces, which suggests that functional mappings between cortical populations are low-dimensional relative to the space of population activity patterns. From this, we partition Fisher information depending on the alignment between the population covariance and the mean tuning direction projected onto the communication subspace or its orthogonal complement. We provide mathematical and numerical analyses of our proposed decomposition of Fisher information and examine theoretical scenarios that demonstrate how to leverage communication subspaces for flexible routing and gating of stimulus information. This work will provide researchers investigating interareal communication with a theoretical lens through which to understand sensory information transmission and guide experimental design.

## 1 Introduction

Complex brain functions, such as behavior and perception, depend on interactions among anatomically and functionally distinct brain regions (Seguin et al. 2023). In sensory areas, neural populations represent the external world by transforming sensory input through local computations; the resulting signal is then transmitted to other cortical areas for further processing. For example, in the mammalian visual system, as visual input moves through the anatomical hierarchy, neurons tend to become increasingly selective (“tuned”) to more complex visual features, from orientation tuning in primary visual cortex (V1) to selectivity for conjuctions of oriented features and for texture in mid-level visual areas V2, V4 and selectivity for object features in higher visual areas (Riesenhuber and Poggio 1999; DiCarlo et al. 2012). This increased selectivity is often characterized by relating changes in mean firing rates throughout neural populations to changes in external input.

Other aspects of neural population activity beyond the firing rate are largely ignored in studies of hierarchical sensory processing. One such feature is the variability of neural responses to repeated presentation of the same stimulus. Neural variability is present throughout the brain (Stein et al. 2005; Goris et al. 2014) and can limit a neuron’s representational capacity (Tolhurst et al. 1983; Geisler and Albrecht 1997). In particular, this neural response variability is often shared throughout a neural population (Cohen and Kohn 2011; Panzeri et al. 2022), and the structure of this shared variability determines the amount of sensory information a population can encode (Shadlen and Newsome 1998; Rumyantsev et al. 2020; Averbeck et al. 2006; Moreno-Bote et al. 2014; Kohn, Coen-Cagli, et al. 2016; Kafashan et al. 2021; Kriegeskorte and Wei 2021). Furthermore, this variability might also constrain the transmission of information between populations in different cortical areas (Zylberberg et al. 2017; Huang, Pouget, et al. 2022; Rowland et al. 2023). With the advance of experimental techniques to record from multiple cortical areas simultaneously (Urai et al. 2022; Abdeladim et al. 2023), new theoretical tools are needed to understand the interplay between neural population variability and sensory transformations (Kang and Druckmann 2020; Kass et al. 2023).

To make progress towards this goal, here we consider a scheme for interareal communication proposed in recent work examining functional interactions in multi-area recordings. Researchers have found that the ability of one cortical area^1^ (the “source” or “sender” area) to influence the activity of a downstream cortical area (the “target” or “receiver” area) depends on how well activity patterns in the source area align with specific directions in the space of neural population activity (Kaufman et al. 2014; Semedo, Zandvakili, et al. 2019; Javadzadeh and Hofer 2022). These axes are referred to as the *communication subspace*, as only fluctuations in activity that lie in this subspace get “communicated” to (i.e. are predictive of the activity of) a downstream region. Empirically, the dimensionality of the communication subspace is significantly less than the dimensionality of both the source and target population activity spaces (Semedo, Zandvakili, et al. 2019; Iyer et al. 2021; Srinath, Ruff, et al. 2021). Communication subspaces have thus been suggested as a mechanism for signal multiplexing, in which multiple different signals can be simultaneously represented by a single neural population and sent to different downstream target areas (MacDowell et al. 2023; Barreiro et al. 2024), and as a means of flexible information routing, in which information is selectively communicated to downstream areas in a context-dependent manner (Kohn, Jasper, et al. 2020; Srinath, Ruff, et al. 2021; Srinath, Ni, et al. 2024; Ritz and Shenhav 2024).

Leveraging this scheme for interareal communication, in this paper we introduce a mathematical framework for studying in detail the transmission of sensory information across neural populations. We focus on the transmission of stimulus-related information between two brain areas. This is different from most studies of functional interactions, including existing work on the subspace, which do not distinguish explicitly between transmission of signals that convey stimulus information and other unrelated shared factors (e.g., endogenous global state fluctuations, Seguin et al. 2023). By examining stimulus-related information transfer, we can concentrate on how the content of a signal (i.e., the stimulus) is transmitted via communication subspaces instead of examining how much of the signal is shared between two areas, regardless of what information it contains (Ince et al. 2015; Celotto et al. 2023). Our method will combine communication subspaces with Fisher information, a widely used measure of stimulus-related information in a neural population that depends on the alignment between the trial-averaged population activity across stimulus values (the signal axis) and the population covariance (Beck et al. 2011; Moreno-Bote et al. 2014).

We aim to decompose the Fisher information in the source and target areas depending on how the neural population activity aligns with the communication subspace. This will allow us to investigate various aspects of communication separately, such as how information is shared between these areas, how information is transmitted from the source area to the target area, and how the received information affects the target area. We provide extensive mathematical derivations of this decomposition and how components of the Fisher information in each area relate to Fisher information lying in different subspaces. To illustrate the explanatory power of our framework, we will then demonstrate how flexible communication of sensory information can be achieved by modifying activity in the source or target areas to change the alignment of population activity to different subspaces of the population activity space.

Our results will guide researchers interested in understanding how stimulus-related information is propagated through the brain via communication subspaces. Because Fisher information (Kanitscheider et al. 2015) and the low-rank linear map (Semedo, Zandvakili, et al. 2019) can be efficiently estimated from neural population recordings with limited data, this method of decomposing Fisher information can easily be used as a data-analytic technique for explicitly measuring stimulus information transmitted through the communication subspace.

## 2 Methods

### 2.1 Defining Communication Subspaces via Reduced-Rank Regression

Consider the combined activity (e.g., spike counts) of all neurons in the source area and arrange them into a population activity matrix 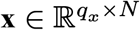, such that the *i*-th column contains the activity of neuron *i* across *N* identical experimental trials. We similarly define the target population activity matrix 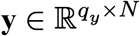. We assume that the source area activity is normally distributed across trials:

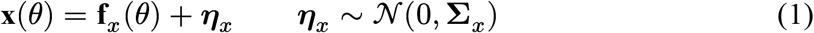

With **Σ**_*x*_ representing the cross-trial noise covariance in the source area and **f**_*x*_ the tuning curve with respect to the stimulus parameter *θ*, which describes how the mean neural activity across experimental trials changes with stimulus. We will assume that all noise covariance matrices are stimulus-independent.

Reduced-rank regression searches for a low-rank matrix (for a prespecified rank *k*) that best approximates the full-rank linear regression matrix of source activity **x** on target activity **y** (Izenman 1975; Tso 1981; Davies and Tso 1982). This can be written as:

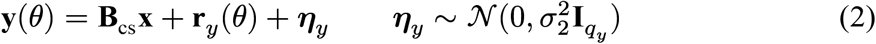

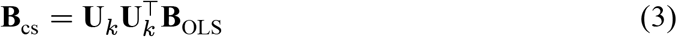

Where **r**_*y*_(*θ*) is the residual tuning in the target area not explained by tuning from the source area (i.e., the tuning curve in the target area is defined as **f**_*y*_ = **B**_cs_**f**_*x*_ + **r**_*y*_), **B**_OLS_ is the full-rank linear regression matrix of **x** on **y, U**_*k*_ are the top-*k* principal components of **B**_OLS_**x** (equivalently, these are the top *k* left singular vectors of 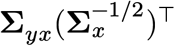, with **Σ**_*yx*_ the cross covariance between **y** and **x**, and 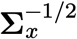 is the inverse Cholesky factor of **Σ**_*x*_, see Izenman 2008, Section 6.3.2), and 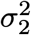 is the residual variance that is minimized in the reduced-rank regression. With this, the noise covariance in the target population, **Σ**_*y*_, is

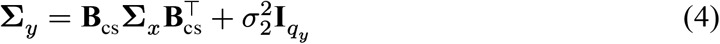

Activity patterns in the source population (i.e., linear combinations of the columns of **x**) that are not communicated to the downstream target population are said to lie in the source population’s private subspace. In other words, an activity pattern υ lies in the private subspace if **B**_cs_υ = 0. Therefore, the private subspace is equivalent to the kernel or null space of **B**_cs_ (written ker(**B**_cs_), which is how we will define the private subspace). Since the vector spaces we are considering in this paper are finite-dimensional Euclidean spaces endowed with the standard inner product, we can then define the communication subspace as the orthogonal complement of the private subspace, ker(**B**_cs_)^⊥^, which is equivalent to the row space of **B**_cs_, Im 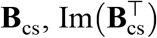 Intuitively, the elements of the communication subspace are those activity patterns that are communicated via the reduced-rank regression mapping **B**_cs_, as vectors in this subspace correspond one-to-one with vectors in Im(**B**_cs_) (Axler 2024, Proposition 6.67).

For Section 2.2, we will want to consider projections onto the communication or private subspaces in the source area. Note that once we define one projection, say the projection onto the communication subspace, which we denote **P**_cs_, then the projection onto the private subspace, **Q**_cs_, is **I** − **P**_cs_. To define these projections, we will use the (Moore-Penrose) pseudoinverse (written **A**^+^), which is a generalization of the inverse of a matrix or linear map to non-invertible transformations (Penrose 1955). As detailed in Appendix A, we can define the projection onto the communication subspace as

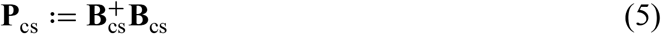

In the preceding discussion, we have implicitly assumed that the reduced-rank regression matrix and all its derived quantities (i.e., the communication and private subspaces and their associated projections) are independent of the stimulus parameter *θ*. We justified this assumption because the closed-form for the reduced-rank regression matrix is a function of the noise covariance matrices (Izenman 1975), which we assumed were stimulus-independent. When applying our method to experimental data, even in the presence of stimulus-dependent covariance, we expect that this will be a reasonable approximation when considering small differences in the stimulus parameter. More generally, researchers should consider if invariant subspaces are a good approximation for their dataset.

### 2.2 Fisher Information Decomposition

The main quantities derived in this section, and their meaning, are summarized in Table 1.

**Table 1.** Table containing definitions of the relevant Fisher information-theoretic terms.

| Symbol | Meaning | Definition (Reference) |
| --- | --- | --- |
| $\mathbf{J}(\mathbf{x}; \theta)$ | Source Fisher Information | $\mathbf{df}_x^\top \Sigma_x^{-1} \mathbf{df}_x$ (Eq 12) |
| $\mathbf{J}(\mathbf{P}_{cs} \mathbf{x}; \theta)$ | Communicated Information | $\mathbf{df}_x^\top \mathbf{P}_{cs} (\mathbf{P}_{cs} \Sigma_x \mathbf{P}_{cs})^+ \mathbf{P}_{cs} \mathbf{df}_x$ (Eq 7, Fig 2A) |
| $\mathbf{J}(\mathbf{Q}_{cs} \mathbf{x}; \theta)$ | Private Information | $\mathbf{df}_x^\top \mathbf{Q}_{cs} (\mathbf{Q}_{cs} \Sigma_x \mathbf{Q}_{cs})^+ \mathbf{Q}_{cs} \mathbf{df}_x$ (Eq 10) |
| $\mathbf{CI}_{cs}$ | Contributed Information to Communication Subspace | $\mathbf{df}_x^\top \mathbf{P}_{cs} \Sigma_x^{-1} \mathbf{P}_{cs} \mathbf{df}_x$ (Eq 14, Fig 2B) |
| $\mathbf{CI}_{priv}$ | Contributed Information to Private Subspace | $\mathbf{df}_x^\top \mathbf{Q}_{cs} \Sigma_x^{-1} \mathbf{Q}_{cs} \mathbf{df}_x$ (Eq 15, Fig 2C) |
| $\mathbf{SI}_1$ | Shared Information between Communication and Private Subspaces | $2\mathbf{df}_x^\top \mathbf{P}_{cs} \Sigma_x^{-1} \mathbf{Q}_{cs} \mathbf{df}_x$ (Fig 3) |
| $\mathbf{J}(\mathbf{y}; \theta)$ | Target Fisher Information | $\mathbf{df}_y^\top \Sigma_y^{-1} \mathbf{df}_y$ (Eq 13) |
| $\mathbf{J}(\mathbf{B}_{cs} \mathbf{x}; \theta)$ | Mapped Information | $\mathbf{df}_x^\top \mathbf{B}_{cs}^\top (\mathbf{B}_{cs} \Sigma_x \mathbf{B}_{cs}^\top)^+ \mathbf{B}_{cs} \mathbf{df}_x$ (Eq 8, Fig 2A) |
| $\mathbf{J}(\mathbf{B}_{cs} \mathbf{x} + \boldsymbol{\eta}_y; \theta)$ | Impactful Information | $\mathbf{df}_x^\top \mathbf{B}_{cs}^\top \Sigma_y^{-1} \mathbf{B}_{cs} \mathbf{df}_x$ |
| $\mathbf{RI}$ | Residual Information | $\mathbf{dr}_y^\top \Sigma_y^{-1} \mathbf{dr}_y$ |
| $\mathbf{SI}_2$ | Synergistic/Redundant Information | $2\mathbf{df}_x^\top \mathbf{B}_{cs}^\top \Sigma_y^{-1} \mathbf{dr}_y$ |

The linear Fisher information **J**(**z**; *θ*) computed from neural activity **z** with tuning curve **f** and noise covariance matrix **Σ** is written as (Kanitscheider et al. 2015):

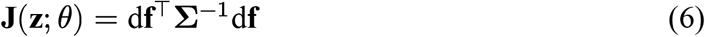

Where d**f** is the derivative of the tuning curve, which is computed between two conditions using the finite difference quotient (i.e.,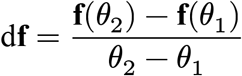), for *θ*_1_ and *θ*_2_ the two different stimulus conditions). This also defines the signal axis in population activity space (Panzeri et al. 2022). The linear Fisher information represents the smallest change in the stimulus that can be discriminated from a neural population using an optimal linear decoder (Kohn, Coen-Cagli, et al. 2016). This is a component of the full Fisher information; in this paper, we will exclusively focus on the linear Fisher information and thus refer to Eq 6 as “Fisher information”.

#### 2.2.1 Fisher Information in the communication and private subspaces

An important quantity for our goal of studying information transmission between areas, is the Fisher information in the communication subspace. There are two possible definitions for this quantity. The first is the Fisher information present in the source population projected into the communication subspace, namely

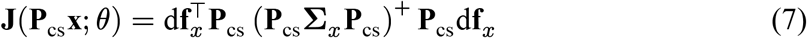

This quantity is closely related to a similar quantity that measures a binary discrimination threshold (*d*^′^)^2^ after using a targeted dimensionality reduction technique (Ebrahimi et al. 2022). The other definition is the Fisher information present in the activity patterns mapped by the reduced-rank regression matrix **B**_cs_:

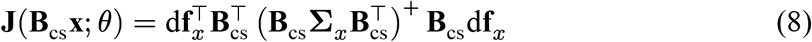

In both expressions, we have to use the pseudoinverse rather than the full matrix inverse, as both matrices are rank-deficient (**P**_cs_**Σ**_*x*_**P**_cs_ is a *q*_*x*_ × *q*_*x*_ matrix of rank *k* while 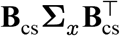 is a *q*_*y*_ × *q*_*y*_ matrix of the same rank).

These two definitions should be equivalent, as **B**_cs_, restricted to the communication subspace, is an isomorphism onto its range, and a simple calculation shows that invertible linear transformations of activity will not change the amount of Fisher information (Huang, Pouget, et al. 2022). However, that calculation does not necessarily translate once we move from matrix inverses into pseudoinverses because, in general, the pseudoinverse of a product of matrices is not the product of their pseudoinverses (Hartwig 1986). Nonetheless, we can prove that these two expressions are equivalent (Appendix B). Thus, we have a well-defined expression for the amount of Fisher information in the communication subspace, which we will denote **J**_cs_:

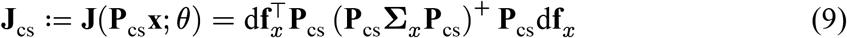

We can similarly define the information in the private subspace as:

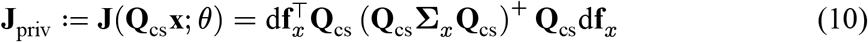

By the data processing inequality for (full) Fisher information (Zamir 1998), we have that the (linear) Fisher information in the source population is an upper bound on the information contained in the communication and private subspaces (see also Appendix B, Lemma 2 which provides a more direct proof of these inequalities as a consequence):

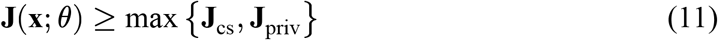

#### 2.2.2 Decomposing Fisher Information in the source and target areas

Using the identity **f**_*x*_ = **P**_cs_**f**_*x*_ + **Q**_cs_**f**_*x*_ (by definition, **P**_cs_ + **Q**_cs_ = **I**) and Eq 6, it is easy to derive the following decomposition of the source Fisher information:

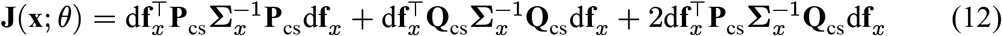

We can similarly decompose the target Fisher information using the identity **f**_*y*_ = **B**_cs_**f**_*x*_ + **r**_*y*_:

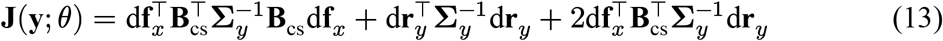

We now provide an interpretation of the different terms in the decompositions above. We will begin with the source Fisher information decomposition and consider the first two terms, which we will define as:

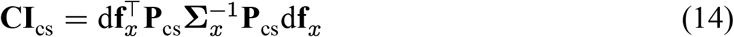

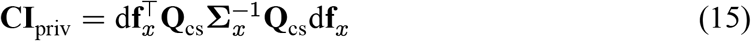

In Appendix C we show (using a technique similar to Zhang 2011, Eq 7.11), that:

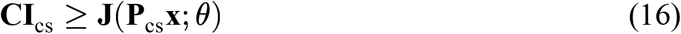

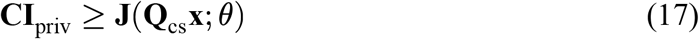

Our understanding of Eqs 14 and 15 is that they are the Fisher information *contributed* to either the communication (Eq 14) or private (Eq 15) subspaces, as they are the Fisher information for a neural population whose tuning curve lies entirely in a subspace (i.e., if **P**_cs_**f**_*x*_ = **f**_*x*_, then the other terms in Eq 12 are 0 and **J**(**x**; *θ*) = **CI**_cs_, and similarly for the private subspace). It is tempting to think that these contributed information terms are themselves bounded by the total source population Fisher information (for example, **J**(**x**; *θ*) ≥ **CI**_cs_). However, this is not true in general as we show in Appendix D.

The final term of the source Fisher information decomposition (Eq 12) is a cross-term that measures the angle between **P**_cs_d**f**_*x*_ and **Q**_cs_d**f**_*x*_ relative to the overall covariance in the source area **Σ**_*x*_. Mathematically, this accounts for the fact that contributed information to the communication and private subspaces can exceed the overall source Fisher information; this cross-term can be thought of as correcting for overcounting when adding these contributed information quantities. Although the communication and private subspaces are orthogonal, that does not mean they are independent (Semedo, Gokcen, et al. 2020). In particular, the information in these subspaces can overlap, as activity in the communication subspace is predictive of the activity in the private subspace and vice versa (Semedo, Zandvakili, et al. 2019). For this reason, we will call this term the *shared information* between the communication and private subspaces:

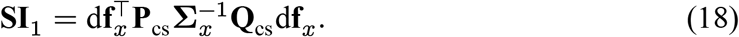

Next, we will consider the components of the target Fisher information decomposition (Eq 13). The first term, which we will call the *impactful information*, is the amount of Fisher information in the target population when **r**_*y*_ = 0, which is when the mean tuning of the source activity completely determines the tuning of the target population. We will denote this term as **J**(**B**_cs_**x** + ***η***_*y*_; *θ*), which is similar in structure to the Fisher information that lies in the communication subspace (Eq 7), except that **J**(**B**_cs_**x** + ***η***_*y*_; *θ*) includes covariance arising from the additional noise ***η***_*y*_ (Eq 4). These terms are related by:

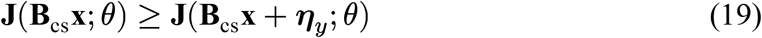

While reduced-rank regression requires that the target noise be a diagonal matrix with isotropic noise (i.e.,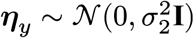), see Eq 2), the above relation also holds if ***η***_*y*_ has an arbitrary covariance matrix **Σ**_*r*_. This is important for possible extensions to the methods proposed here that would apply to related dimensionality reduction techniques such as canonical correlation analysis that have been used in the literature to characterize cortical communication (Semedo, Jasper, et al. 2022; Semedo, Gokcen, et al. 2020; Gokcen, Jasper, Xu, et al. 2024). There are two ways to demonstrate that this relationship holds. Indirectly, this can be seen as another consequence of the data processing inequality for Fisher information (Zamir 1998), as the chain relationship *θ* → **B**_cs_**x** → **B**_cs_**x** + ***η***_*y*_ is trivially a Markov chain. The more direct argument is provided in Appendix E.

The second term in the target Fisher information (Eq 13) will be called the *residual information* (**RI**). This term would reflect the stimulus-related information present in the target area when no tuning information is linearly communicated between the source and target (i.e., when 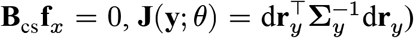) or equivalently, after subtracting the influence of the source area on the target.

Lastly, the final term in the target Fisher information decomposition (Eq 13) provides a measure of the interaction between the impactful information and the residual information, as it measures the angle between **B**_cs_d**f**_*x*_ and d**r**_*y*_ relative to principal axes of the target covariance matrix **Σ**_*y*_. For this reason, we will call it the *synergistic/redundant information* between the residual and transmitted tuning:

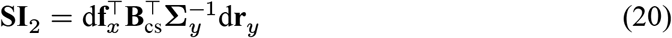

This paper will not examine these terms as we assume no residual tuning for simplicity (see Section 2.3).

#### 2.2.3 Considerations on estimation from finite data

With our choice to focus on Fisher information, applications of the framework proposed here to real data can leverage existing methods to estimate linear Fisher information efficiently and accurately for large neural populations. These methods include linear decoding (Kohn, Coen-Cagli, et al. 2016) and direct bias-corrected estimators (Kanitscheider et al. 2015). The direct estimators are straightforward; they require measuring the empirical tuning derivatives (by finite difference) and noise covariances, projecting them through the appropriate mapping matrices, and applying analytical bias corrections. Using linear decoding requires some additional considerations. Specifically, some terms of our decompositions do not immediately correspond to decoding measured activity. For instance, **CI**_cs_ uses the full noise covariance of the source area, but only the tuning derivatives projected onto the subspace (not the full tuning of the source area, see Eq 14), so it cannot be estimated by decoding the source area activity. However, in this case, it is easy to construct a transformation of the target area activity **x** − **Q**_cs_**f**_*x*_, which has exactly those tuning derivatives and covariance. Therefore, **CI**_cs_ can be estimated by decoding after applying the transformation above. Similarly, the impactful information term can be estimated with a linear decoder computed from **y** − **f**_*y*_ + **B**_cs_**f**_*x*_. Lastly, the term denoted **SI**_1_ can be computed from data by using the linear decoders for **x, CI**_cs_ and **CI**_priv_ (discussed above) with Eq 12 and, similarly, we can also estimate **SI**_2_ from data by combining estimates for **J**(**y**; *θ*), **J**(**B**_cs_**x** + ***η***_*y*_; *θ*) and **RI**.

### 2.3 Simulations

In Section 3, we will analyze Fisher information and derived quantities (the quantities in Eqs 9, and 10, 12, and 13) computed directly from simulated tuning curves, reduced-rank regression matrices, and covariance matrices.For the baseline simulations performed, we need to specify the tuning curves **f**_*x*_, **f**_*y*_, the source covariance **Σ**_*x*_, the reduced-rank regression matrix **B**_cs_, and the residual variance 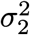 The number of neurons in the source (*q*_*x*_) and target (*q*_*y*_) area will be 50. We will assume that there is no residual tuning in the target area, such that all the stimulus-related information in the target area arises from the feedforward contribution of the source area (**f**_*y*_ = **B**_cs_**f**_*x*_). The tuning in the source area will be von-Mises tuning curves, which resemble the orientation selectivity of neurons in V1.The stimulus parameter, *θ*, will take two values: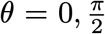. The *i*-th component of **f**_*x*_, *f*_*x,i*_ is parameterized as

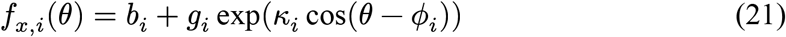

Where *b*_*i*_ is the bias, *g*_*i*_ is the amplitude, *ϕ*_*i*_ is the preferred orientation for neuron *i*, and *κ*_*i*_ controls the width of the tuning curve. Across all neurons in the source area, *b*_*i*_ = 2, *g*_*i*_ = 30/*e*. The value of *ϕ*_*i*_ is drawn from 𝒰([0, *π*]), while the value for *κ*_*i*_ is drawn from 𝒰([0.5, 2]). The correlation structure is generated as follows: first, we generate a limited range correlation matrix **R**_*ϕ*_ from the preferred orientations *ϕ*_*i*_ (with spatial scale *L* = 1 and correlation scale *c*_0_ = 0.3, see Ecker et al. 2011). Then we draw a random correlation matrix **R**_*LKJ*_ from LKJ distribution with *β* parameter fixed at 30 (Lewandowski et al. 2009) and set a mixing proportion *s* = 0.8 to create the source correlation matrix **R**_1_ = *s***R**_*ϕ*_ +(1 −*s*)**R**_*LKJ*_. This allows for some negative noise correlations between neurons, as all limited-range correlations are positive. To form the noise covariance matrix **Σ**_*x*_, we set the variances to be proportional to the mean (“Poisson-like variability”): for all neurons *i*, we took the mean of the tuning curve across the two stimulus values 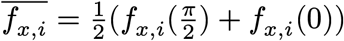, created a diagonal standard deviation matrix **S** with diagonal entries 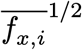, and set **Σ**_*x*_ = **SR**_1_**S**. The reduced-rank regression matrix is generated by taking a truncated SVD of a random Gaussian matrix: we generate a *q*_*y*_× *q*_*x*_ matrix **B** with entries sampled from 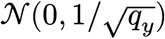 compute the SVD **B** = **USV**^⊤^, and retain only the top *k* singular values and vectors. We fixed the dimensionality of the communication subspace (the number of singular values to retain during truncation) at *k* = 5, on par with empirical data (Semedo, Zandvakili, et al. 2019). Finally, we fix the residual variance 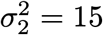.

Additionally, we will modify the structure of the tuning curves and covariance matrices to exemplify certain properties of the information decomposition we are proposing and how this would influence using communication subspaces as a flexible stimulus-related information propagation scheme. These manipulations will be done to assess how manipulations of the model structure affect Fisher information quantities. This allows direct comparisons (per simulation) of the information quantities computed from the initially generated parameters and their perturbations.

## 3 Results

The communication subspace framework explains fluctuations in activity in a target neural population from a low-dimensional subset of activity in a source population (Fig 1). Intuitively, only information in this predictive subspace (the communication subspace) should be inherited by the activity in the target area. In contrast, information in the orthogonal complement (the private subspace) is not communicated to this area and can be used for local processing or communicated to a different target area (Semedo, Zandvakili, et al. 2019). The linkage between the two areas, the communication subspace, is analogous to a bottleneck through which information present in the source area must flow to get to the target area (although this is distinct from the information bottleneck technique proposed by Tishby et al. 2000). This paper examines these concepts in a more rigorous setting by investigating how stimulus-related information is transmitted through this system using Fisher information (Eq 6), which measures how well the stimulus values can be linearly decoded from neural population activity.

**Figure 1.**
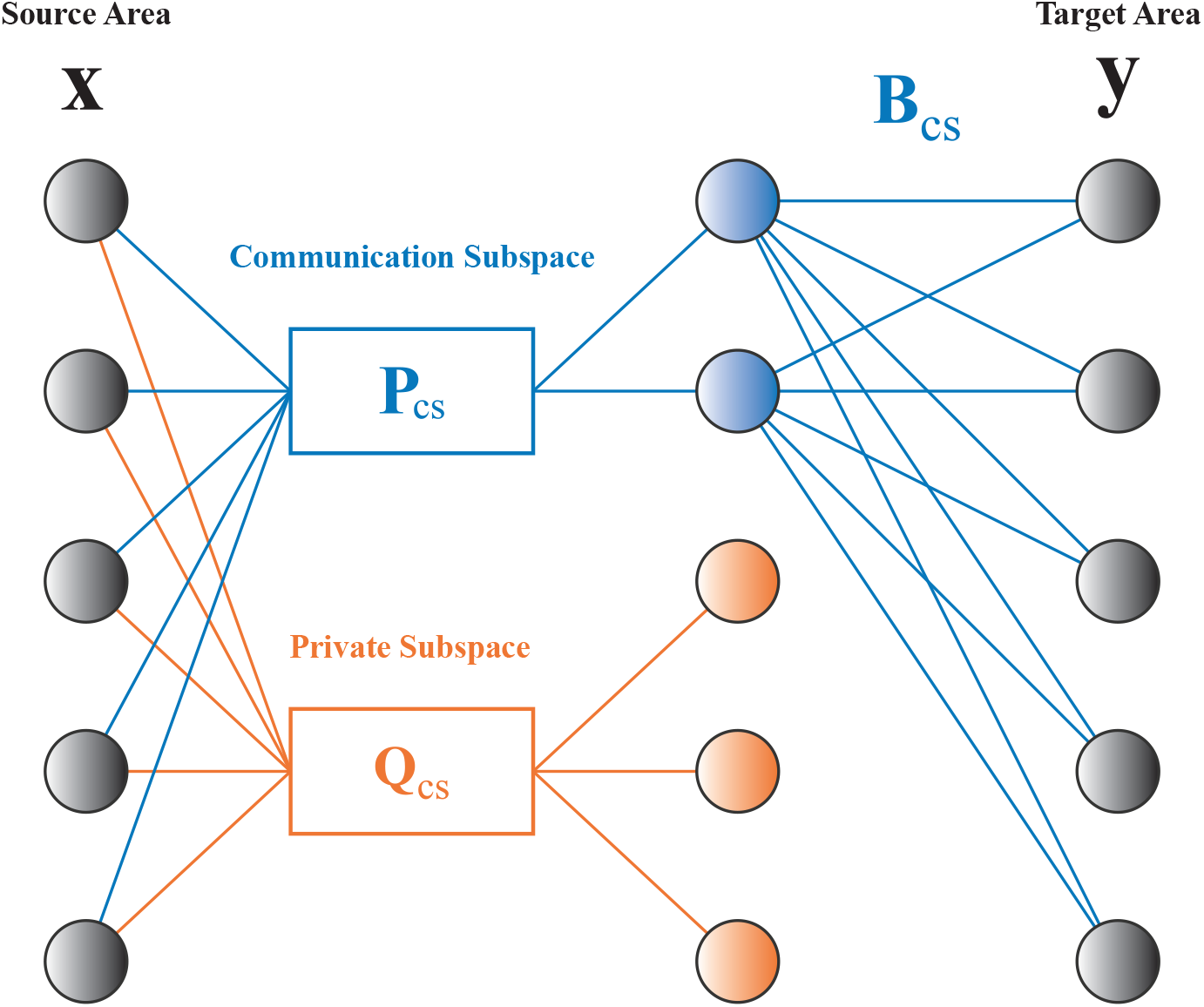
Communication Subspace. Representation of communication between two cortical areas (source and target) as a linear autoencoder. Linear combinations of neural activity in the source area can be divided into two orthogonal subspaces: the communication subspace (blue) and the private subspace (orange). Only activity in the communication subspace is used to linearly predict activity in the target area through the map **B**_cs_. The activity in the private subspace lies in the kernel of **B**_cs_, so is mapped to 0.

### 3.1 Measuring stimulus-releated information in communication subspaces

We first illustrate our proposed framework to characterize stimulus-related information transmission through the communication subspace, using linear Fisher Information **J** about a stimulus variable *θ* (details and proofs in Section 2.2; see also Fig 1 and Table 1 for definitions of terms). There are two possible definitions for the information “in” the communication subspace: the information computed from the source activity projected onto the communication subspace (**J**(**P**_cs_**x**; *θ*), Eq 7) and the information computed from the source activity mapped ono the target via the reduced-rank regression matrix (**J**(**B**_cs_**x**; *θ*), Eq 8). We showed analytically and with simulations (Fig 2A) that these quantities are equal (the common term written **J**_cs_, Eq 7). Similarly, we also defined the term **J**_priv_ (Eq 10) denoting Fisher information in the private subspace.

**Figure 2.**
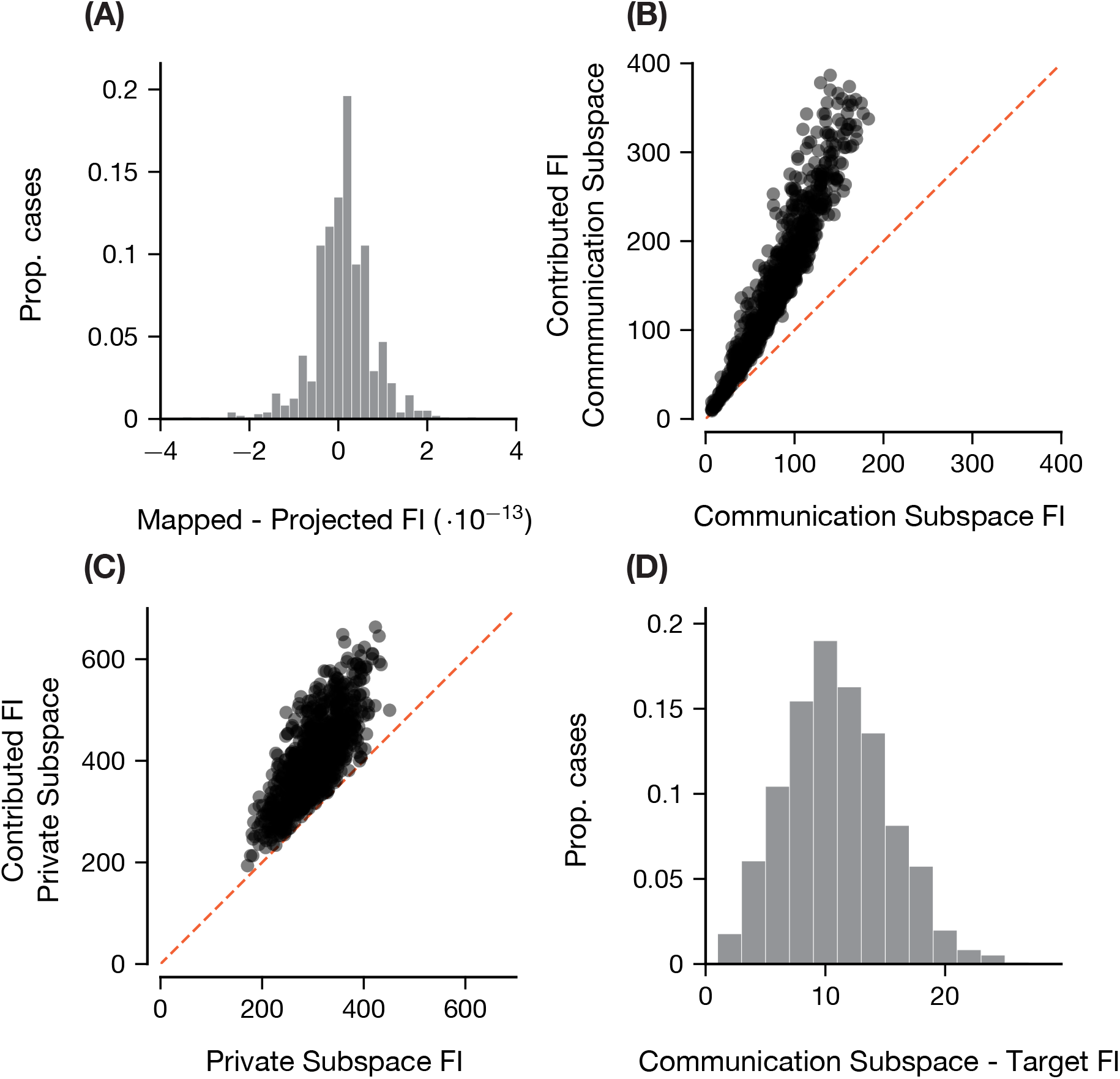
Validation of Relationships between Fisher Information Theoretic Terms. Simulated population means and covariance matrices were generated as described in Section 2.3. Fisher information theoretic terms were computed for each simulation. 1000 simulated mean vectors and noise covariance matrices were generated. (A) Information in the communication subspace **J**_cs_ has a consistent definition (Eq 30): **J**(**B**_cs_**x**; *θ*) (Eq 8) = **J**(**P**_cs_**x**; *θ*) (Eq 7) (B) Contributed information to the communication subspace is an upper bound on information in the communication subspace (Eq 16): **CI**_cs_ (Eq 14)≥ **J**_cs_(C) Contributed information to the private subspace is an upper bound on information in the private subspace (Eq 17): **CI**_priv_ (Eq 15) ≥ **J**_priv_ (Eq 10) (D) Information in the communication subspace is an upper bound on the impactful information (information received from the source population, Eq 19): **J**_cs_ ≥ **J**(**B**_cs_**x** + ***η***_*y*_; *θ*). Note that in these simulations the impactful information is equivalent to the Target information because **r**_*y*_ = 0.

Next, we derived equations (Eqs 12 and 13) that decompose information in the source and target (**y**) areas with respect to the low-rank linear mapping between the two areas (Eq 2). In the decomposition for the source area, there are two terms (**CI**_cs_ and **CI**_priv_, Eqs 14 and 15) that are very similar to the definitions for the Fisher information in the communication and private subspaces (**J**_cs_ and **J**_priv_), except that they use the full source area covariance **Σ**_*x*_ as opposed to the covariance of the source activity after being projected into the corresponding subspace. We proved a lemma (Appendix C) which demonstrated that **CI**_cs_ and **CI**_priv_ provide upper bounds for **J**_cs_ and **J**_priv_ (Eqs 16 and 17; simulations in Fig 2B and 2C). For this reason, we named **CI**_cs_ and **CI**_priv_ the *contributed information* to either the communication or private subspace. Lastly, we showed that **J**_cs_ is an upper bound on the information present in the target area when all stimulus-related activity in the target population is entirely driven by the source population (**J**(**B**_cs_**x** + ***η***_*y*_; *θ*), Fig 2D). Therefore, we termed this latter quantity *impactful information* because it measures how much the communicated information impacts the target area.

The Fisher information decomposition in the source area (Eq 12) contains an additional term, **SI**_1_. As described in Section 2.2, this term measures the angle between the projections of the signal axis (d**f**_*x*_) onto the communication and private subspace relative to principal axes of the source population activity space (i.e., the eigenvectors of **Σ**_*x*_). Because of this, we claimed that this term accounts for how much information is shared between the communication and private subspaces, and we called it *shared information*. In particular, as the other terms in the source Fisher information decomposition (the contributed information terms **CI**_cs_ and **CI**_priv_) can be larger than the overall source Fisher information, the shared information term corrects for overcounting when combining these two terms. We tested this idea by measuring how **SI**_1_ changes with the size of the communication subspace (Fig 3A). For low- and high-dimensional communication subspaces (relative to the dimensionality of the source area), the magnitude of the cross-term is relatively small because there is less overlapping information between the communication and private subspaces due to their complementary sizes (e.g., when the communication subspace is low-dimensional the private subspace is high-dimensional, and vice versa). When the communication and private subspaces are of comparable dimensionality, the shared information is highly negative, which is due to the communication and private subspaces containing redundant information. We quantified this by comparing the shared information and the sum of Fisher information in the communication and private subspaces (**J**_cs_ + **J**_priv_, Fig 3B). There is a strong negative correlation (−0.75), indicating that the more overall information in both the communication and private subspace, the more likely this information is not unique to these subspaces. This provides some evidence that the shared information term is responsible for measuring the extent to which the communication and private subspace have overlapping or redundant stimulus-related information.

**Figure 3.**
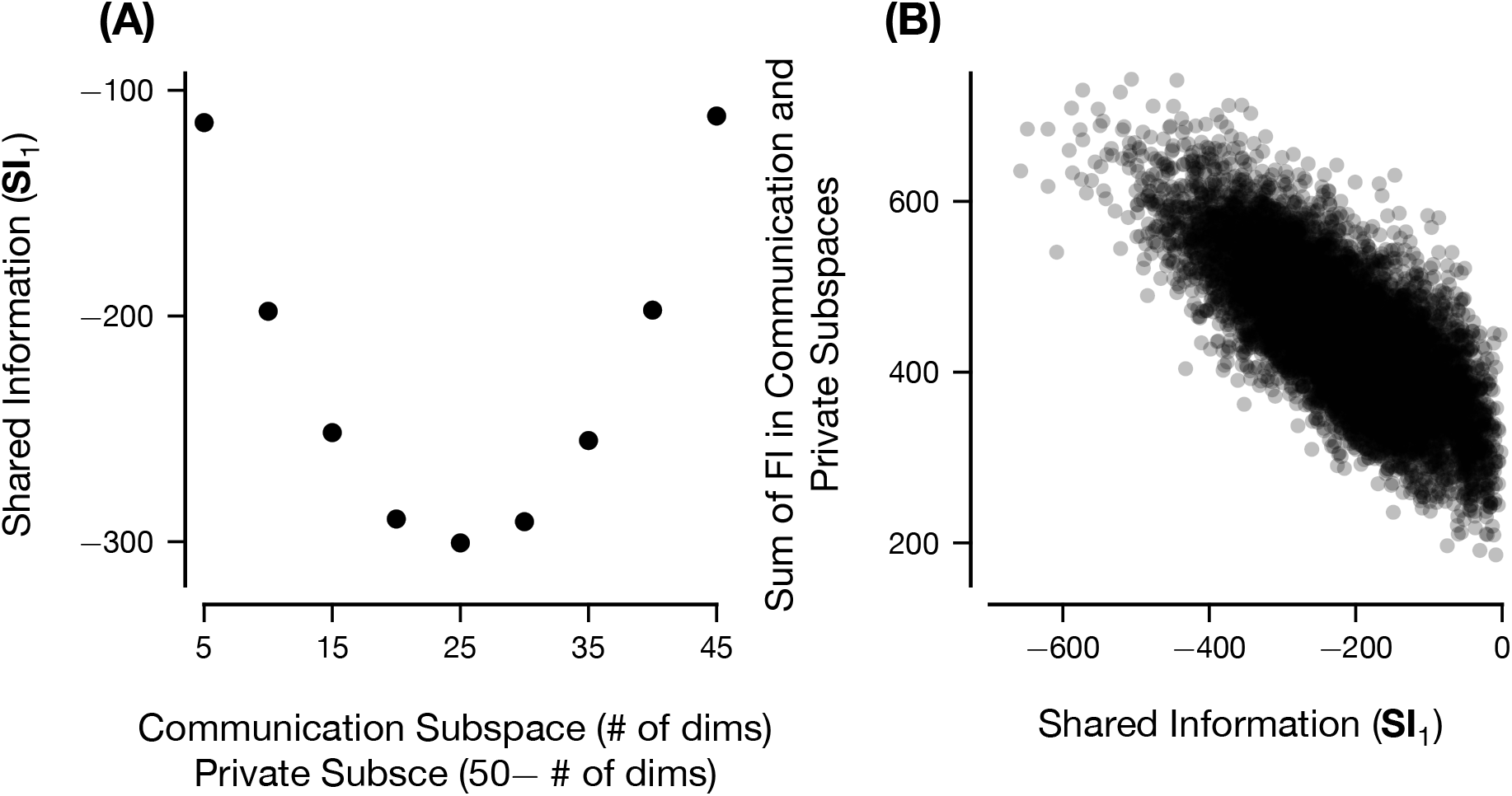
Shared Information between Communication and Private Subspace. (A) The relationship between the mean shared information term in the source area **SI**_1_ and the dimensionality of the communication subspace (or equivalently, 50 (the dimensionality of the source population) − the dimensionality of the private subspace). (B) How the magnitude of the shared information in the source area varies with the sum of Fisher information in the communication subspace and private subspace **J**_cs_ + **J**_priv_. Each point in (A) represents the average across 1000 simulated experiments with different tuning curves and covariance matrices. Dimension of the communication subspace was varied between 5 and 45. Each point in (B) represents a single simulation with fixed number of communication subspace dimensions between 5 and 45.

In summary, we have derived mathematical equations for decomposing Fisher information in the source and target areas with respect to a low-rank linear map that describes the functional connectivity between these areas. We verified the relationships among the key quantities in this decomposition analytically and numerically (Fig 2). We then showed how 1) the impact on the target area (impactful information) does not simply depend on the communicated information, but also on the variability in the target area, with important consequences for the optimality of transmission and, as we show later, for selective gating at the target; 2) one term in the source Fisher information decomposition, the shared information **SI**_1_, describes the extent to which the communication and private subspaces contain the same information about the stimulus (Fig 3). These simulations validate our intuitions (detailed in Section 2.2) regarding the interpretation of the different components of the Fisher information decompositions. Next, we demonstrate a few scenarios in which our theory that can help us better understand information transmission across brain areas.

### 3.2 Selectively Altering Communication of Information by Modifying Tuning

#### 3.2.1 Selectively Increasing Information in the Private Subspace

One of the scenarios presented in the original communication subspaces paper concerns the routing of information to two downstream cortical areas (Semedo, Zandvakili, et al. 2019). In this scenario, two downstream target populations (*T*_1_ and *T*_2_) receive input from a single source area (*S*) through two different reduced-rank regressions, corresponding to two different communication subspaces. How would the source area *S* be able to selectively communicate with downstream target area *T*_1_ without affecting communication between *S* and *T*_2_ and vice versa? Semedo, Zandvakili, et al. propose one solution: if the private subspace for communication between *S* and *T*_2_ is contained in the communication subspace that exists between *S* and *T*_1_ (symbolically, ker(*S* → *T*_2_) ⊂ ker(*S* → *T*_1_)^⊥^), then fluctuations in activity in *S* that lie entirely in ker(*S* → *T*_2_) are communicated to *T*_1_ but do not affect information transfer from *S* → *T*_2_. This alignment between the communication and private subspaces would allow the source area to represent multiple signals simultaneously and transmit different signals to distinct downstream areas (Jun et al. 2022; MacDowell et al. 2023).

We will formalize this observation using our Fisher information decomposition. Consider the case in which one source area communicates with one target area. First, we will demonstrate that one can increase the amount of information in the private subspace without affecting the communication between source and target. Then, conversely, we will consider how to increase information in the communication subspace without affecting information in the private subspace. This is a simplification of the scenario we previously presented, but maintains the core idea: if we can increase information in the private subspace without altering the information in the communication subspace or target area, then a different target area, whose communication subspace contains this private subspace, can receive this information.

To demonstrate this possibility, we first need to show that we can increase information in the private subspace without changing communication to the target area. Suppose a source area with tuning curve **f**_*x*_ is communicating with a target area via the low-rank linear mapping **B**_cs_. Then we can decompose **f**_*x*_ = **P**_cs_**f**_*x*_ + **Q**_cs_**f**_*x*_, where **Q**_cs_ is the projection onto ker(**B**_cs_) (see Section 2.2). Now, we will modify the tuning curve by increasing the projection of the tuning curve onto the private subspace:

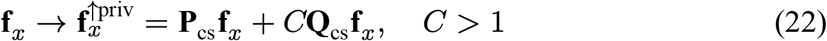

We will call the source activity with this new tuning curve **x**^↑priv^. To begin, we assume that the covariance matrix for the modified source activity is unchanged 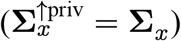.

Fig 4 shows the results of 1000 simulations in which we increased the private tuning (Eq 22 with *C* = 1.5). We can see in Fig 4A that this modification causes the relationship between the source Fisher information and target Fisher information to shift primarily to the right (black is the baseline, orange is after modifying the tuning curve), indicating that we are increasing the source Fisher information without affecting the target Fisher information. This can be seen directly in the following panels in which we compare the source Fisher information (Fig 4B) and target Fisher information (Fig 4C) before and after increasing the private tuning. Clearly, after modifying the private tuning, the source Fisher information increases while the target Fisher information stays the same. In essence, the increase in the source Fisher information is because we are increasing the amount of information in the private subspace (Fig 4D), which we can think of as a component of the source Fisher information. Thus, increasing the information in the private subspace should increase the source Fisher information^2^. Additionally, as we are not changing the amount of information in the communication subspace (Fig 4E), there should be no change in the amount of impactful information **J**(**B**_cs_**x** + ***η***_*y*_; *θ*), which for our simulations is exactly the amount of information in the target area. We also derived these effects mathematically by directly comparing the Fisher information terms (Appendix F).

**Figure 4.**
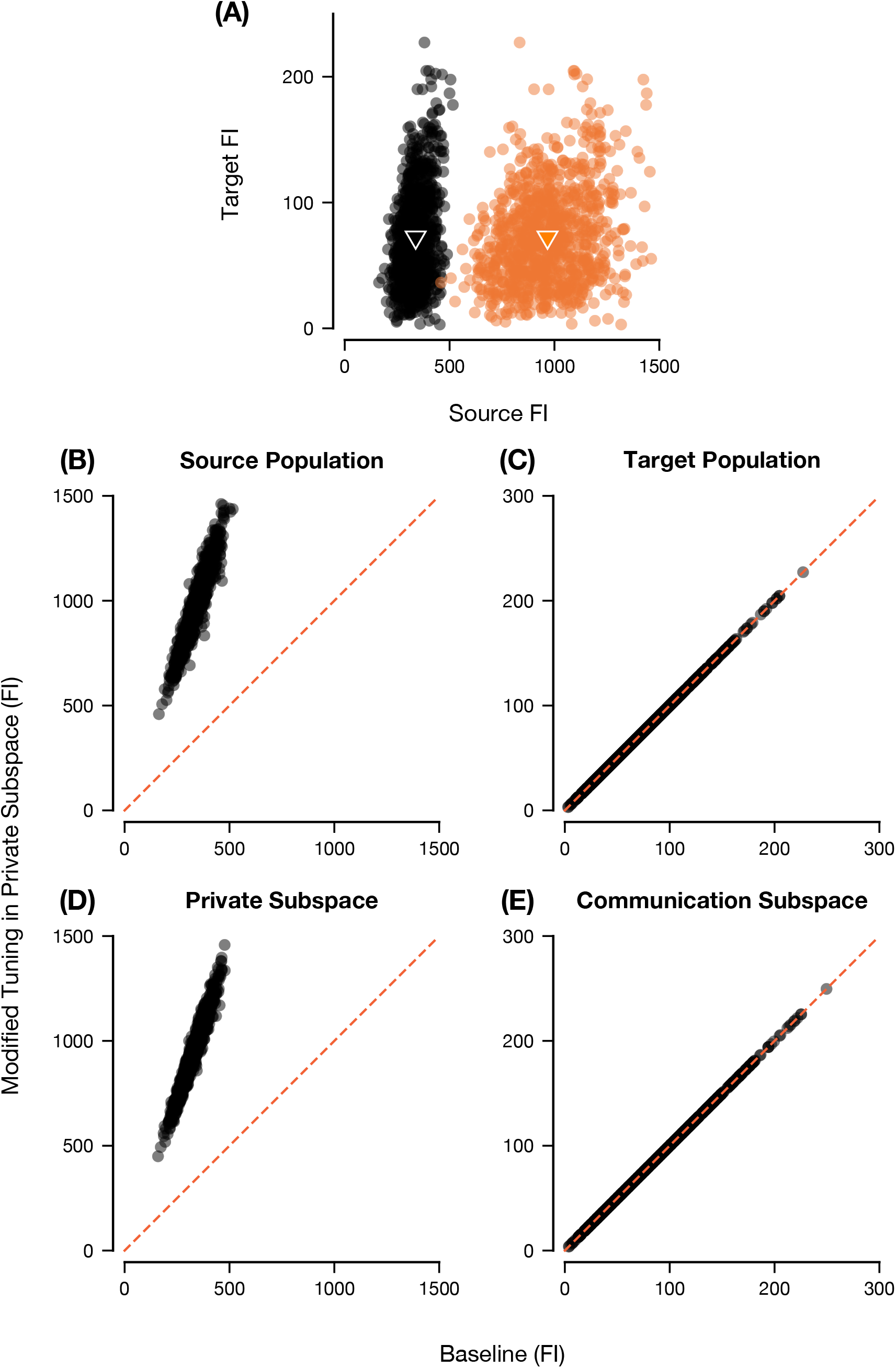
Changes in Fisher Information after Modifying Tuning in the Private Subspace. (A) Scatter plot of Fisher information (FI) in the source area vs. FI in the target area before (black) and after (orange) modifying the private tuning. The triangular marks indicate the means of the respective point clouds. (B) Fisher information in the source area before and after modifying the private tuning. The x-axis corresponds to the abscissa of the black scatter points in (A), while the y-axis corresponds to the abscissa of the orange scatter points in (A). (C) Fisher information in the target area before and after modifying the private tuning. The x-axis corresponds to the ordinate of the black scatter points in (A), while the y-axis corresponds to the ordinate of the orange scatter points in (A). (D-E) Fisher information in the private subspace (D) and communication subspace (E) before and after modifying tuning in the private subspace. 1000 simulated tuning curves and covariance matrices were created. For each simulation, the tuning curve was modified using Eq 22 with *C* = 1.5, with each neuron (i.e., element of the tuning curve vector **f**_*x*_) having some additional random jitter for visualization purposes.

As discussed in Section 2.3, we created the baseline source covariance matrix such that the variance would be equal to the mean. However, by modifying the source area tuning curve, we have broken this power-law relationship that is often observed in neural recordings. To address this, we can also modify the covariance matrix using the same procedure used to generate the source covariance matrix with the new tuning curve (Eq 22) and compare the Fisher information quantities for **x** and **x**^↑priv^ (Fig 5). In this case, many of the same conclusions apply numerically; however, the Fisher information in the communication subspace and the target area decrease (Fig 5C, E). This is likely occurring because, by increasing tuning in the private subspace, we tend to increase the average across-stimulus response of neurons in the source area, which increases the response variance. If this also causes the variability in the communication subspace to increase (i.e., if the eigenvalues of 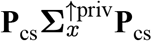 are greater than those for **P**_cs_**Σ**_*x*_**P**_cs_), then the Fisher information quantities computed from the covariance in the communication subspace will tend to decrease. Intuitively, since the Fisher information depends on the (pseudo)inverse of the covariance matrix (Eq 6), by increasing variability in this subspace we are “dividing” by a larger quantity, thus decreasing the information. This is not entirely true, however, as the effect will also depend on how the principal axes of activity in the communication subspace (i.e., the eigenvectors of **P**_cs_**Σ**_*x*_**P**_cs_) align with the projected signal axis **P**_cs_d**f**_*x*_ (Zylberberg et al. 2017, Eq 71) and the size of the subspace being modified; it is possible for the Fisher information to increase after this modification (see Fig 8D and the explanation related to Fig 7).

**Figure 5.**
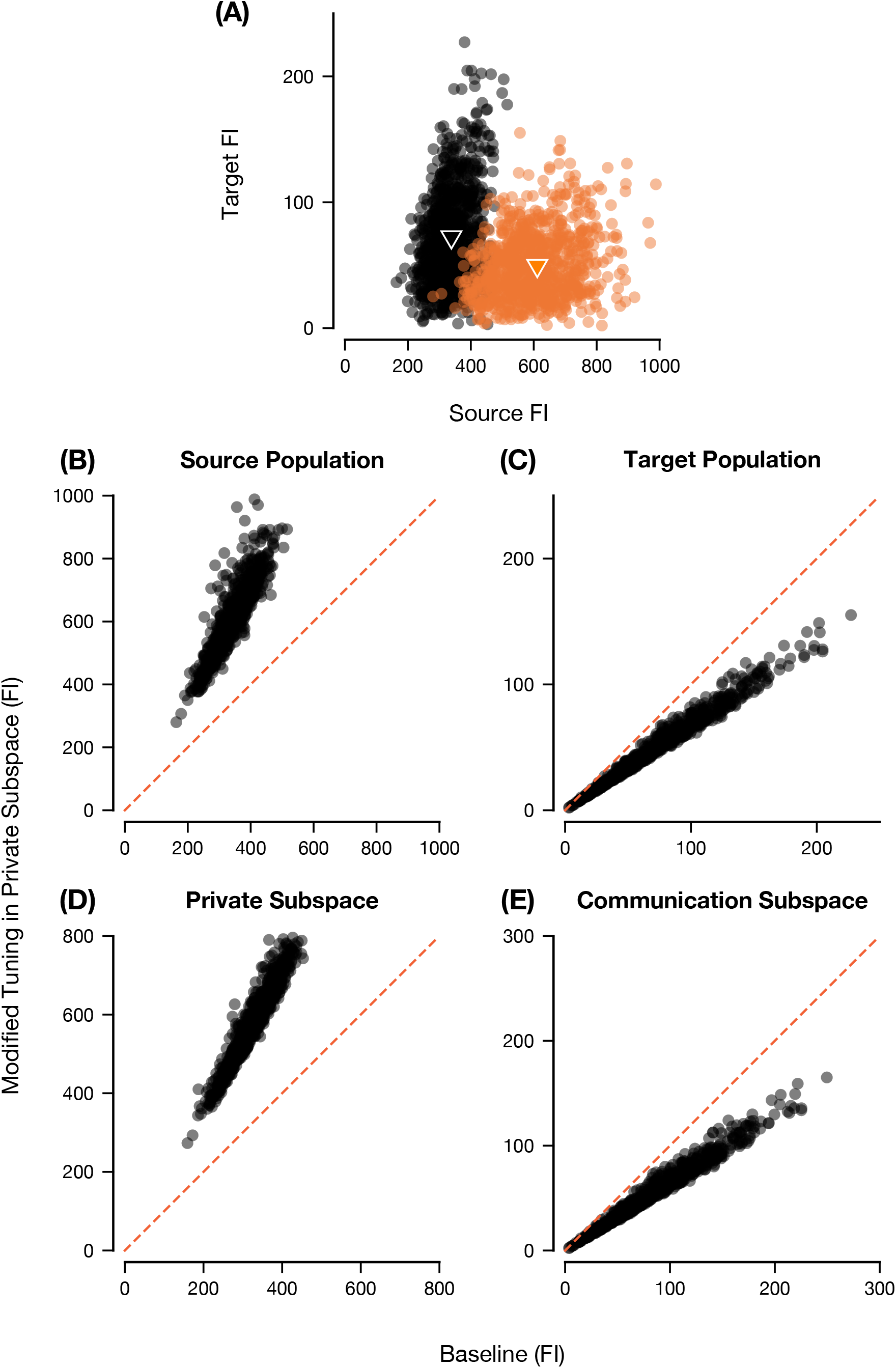
Changes in Fisher Information after Modifying Tuning and Covariance in the Private Subspace. Same as Fig 4 except we modify the covariance matrix so that the variance is equal to the mean.

In summary, the analysis above demonstrates that stimulus information can be increased selectively in one area without affecting all its targets, by changing tuning in the private subspace (Fig 4). However, it also cautions that if the tuning modification is accompanied by changes in noise covariance, then transmission to the target may be impaired (Fig 5), which is an important consideration for realistic circuit models of selective communication (see Section 4).

#### 3.2.2 Selectively Increasing Information in the Communication Subspace

We can use a similar method as in Section 3.2.1 to increase the strength of communication between a source and target area while not altering the amount of information in the private subspace. To do so, we replace the tuning curve with one in which we increase the contribution of the projection onto the communication subspace:

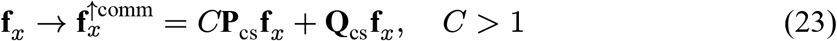

We will refer to the source population activity with this modified tuning curve as **x**^↑comm^ (again note the covariance matrix will initially be unchanged).

Fig 6 is analogous to Fig 4, except we modified tuning in the communication subspace instead of modifying the private tuning (Eq 23 with *C* = 1.5). Before modification, the baseline tuning curves and covariance matrices are the same in Figs 4 and 6. Looking at Fig 6A and comparing the positions of the means (triangular marks) of the point clouds for both the baseline (black) and modified tuning in the communication subspace (blue), it is clear that this modification to the communication subspace causes the relationship between source and target Fisher information to shift up and to the right, indicating an increase in both the source and target areas. The increase in Fisher information in the source and target populations is due to the increase of Fisher information in the communication subspace (Fig 6E, but again note the same caveat applies here as when we considered increasing information in the private subspace). In contrast, the Fisher information in the private subspace does not change (Fig 6D). Mathematically, these effects can all be explained similarly to the change of tuning in the private subspace (Appendix F).

**Figure 6.**
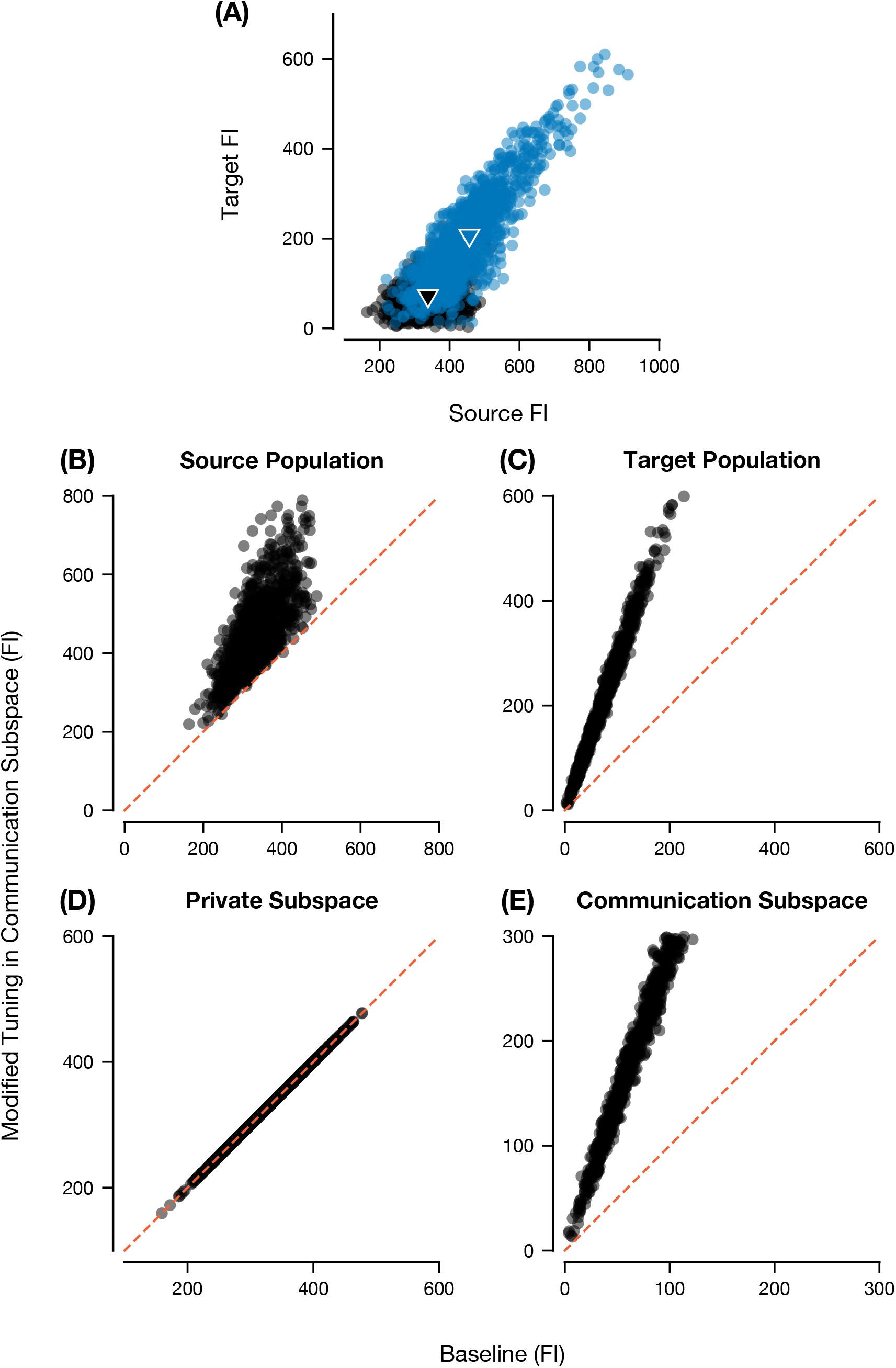
Changes in Fisher Information after Modifying Tuning in Communication Subspace. (A) Scatter plot of Fisher information (FI) in the source area vs. FI in the target area before (black) and after (blue) modifying the tuning in the communication subspace (blue). The triangular marks indicate the means of the respective point clouds. (B) Fisher information in the source area before and after modifying the tuning in the communication subspace. The x-axis corresponds to the abscissa of the black scatter points in (A), while the y-axis corresponds to the abscissa of the blue scatter points in (A). (C) Fisher information in the target area before and after modifying the tuning in the communication subspace. The x-axis corresponds to the ordinate of the black scatter points in (A), while the y-axis corresponds to the ordinate of the blue scatter points in (A). (D-E) Fisher information in the private subspace (D) and communication subspace (E) before and after modifying tuning in the communication subspace. 1000 simulated tuning curves and covariance matrices were created. For each simulation, the tuning curve was modified using Eq 23 with *C* = 1.5, with each neuron (i.e., element of the tuning curve vector **f**_*x*_) having some additional random jitter for visualization purposes.

We also noted that the increase in source area (Fig 6B) is much more modest, especially in comparison to the increase in source area Fisher information when increasing private tuning (Fig 4B). The reason for this is due to the difference in dimensionality between the communication subspace and the private subspace. In our simulations (and in neural data (Semedo, Zandvakili, et al. 2019)), the dimensionality of the communication subspace is significantly less than the dimensionality of the neural population space (the dimension of the communication subspace in our simulations is 5, while the number of neurons in the source population is 50). For this reason, when we increase tuning in the private subspace, it is approximately equal to scaling the whole population mean tuning curve, which causes a noticeable effect on the amount of Fisher information in the source area (and Fisher information contained in the private subspace, Fig 4B and 4D). Conversely, increasing tuning in the communication subspace modifies a small portion of activity in the source area, so the effect of Fisher information in the source area is not as large (Fig 6B and notice the smaller magnitude of information in the communication subspace, Fig 6C). We can see this in Fig 7, in which we increase the dimensionality of the communication subspace and measure how much the Fisher information in the source area increases when we modify tuning in the communication (blue bars) or private (orange bars) subspace. Clearly, as the dimensionality of the communication subspace increases, the Fisher information in the source area after modifying tuning in the communication subspace increases (relative to the unmodified baseline), while the same monotonic effect is observed for modifying tuning in the private subspace. This also explains the difference in scale between the source and target Fisher information quantities in general, as only the stimulus-related changes lying in the low-dimensional communication subspace are transmitted to the target area (Figs 4A and 6A).

**Figure 7.**
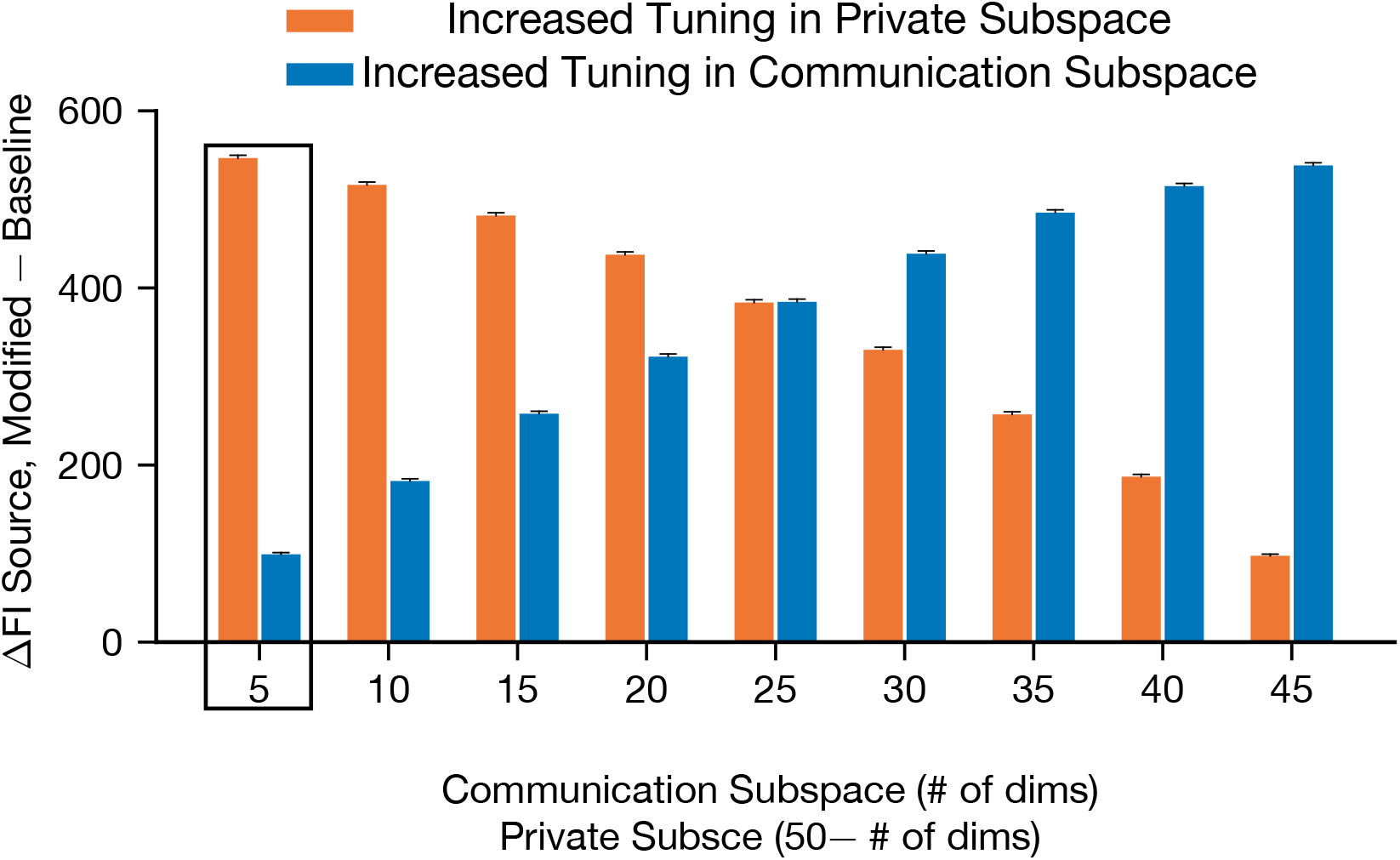
Increase in Fisher Information in Source Area after Modifying Tuning Scales with Size of Subspace. Difference between Fisher information in the source area with increased tuning in either the private subspace (orange) or communication subspace (blue) to Fisher information in the source area at baseline (**J**(**P**_cs_**x**^↑comm or priv^; *θ*) − **J**(**P**_cs_**x**; *θ*)). Each bar represents the average difference across 1000 simulations with the dimensionality of the communication subspace fixed for a value between 5 (which corresponds to the simulations in Figs 4 and 5 (orange bar), and Figs 6 and 8 (blue bar)) and 45. By construction, the dimensionality of the private subspace for each simulation is 50 (the number of neurons in the source area) − the dimensionality of the communication subspace. As with Fig 6, the tuning curves were modified using Eq 23 with *C* = 1.5 with some per-neuron jitter. Error bars are the standard error of the mean difference.

If we modify both tuning curves and covariance covariance matrix to maintain the powerlaw relationship between the variance and the mean, we find qualitatively similar effects (Fig 8). The only noticable difference is for the Fisher information in the private subspace; for some simulations, the Fisher information increases after modifying the tuning curve in the communication subspace and the covariance matrix, while for others it decreases (Fig 8D). However, for the vast majority of cases, these differences were not significant (934/1000 simulations had absolute value of the z-scored difference ≤ 1.96). The reason there is a stronger effect on the Fisher information in the communication subspace after increasing private tuning (Fig 5E) than for the Fisher information in the private subspace here (Fig 8D) is likely also due to the dimensionality of these subspaces in our simulations. As explained previously, for our simulation, increasing tuning in the private subspace should nearly increase tuning for the whole space, so the variance of all neurons in the source area will increase. Increasing variability in all directions will tend to also increase variability in the communication subspace, so the Fisher information should decrease. On the other hand, because the effect of increasing tuning in the communication subspace does not greatly effect the tuning curve magnitude, the covariance matrix will not change substantially.

**Figure 8.**
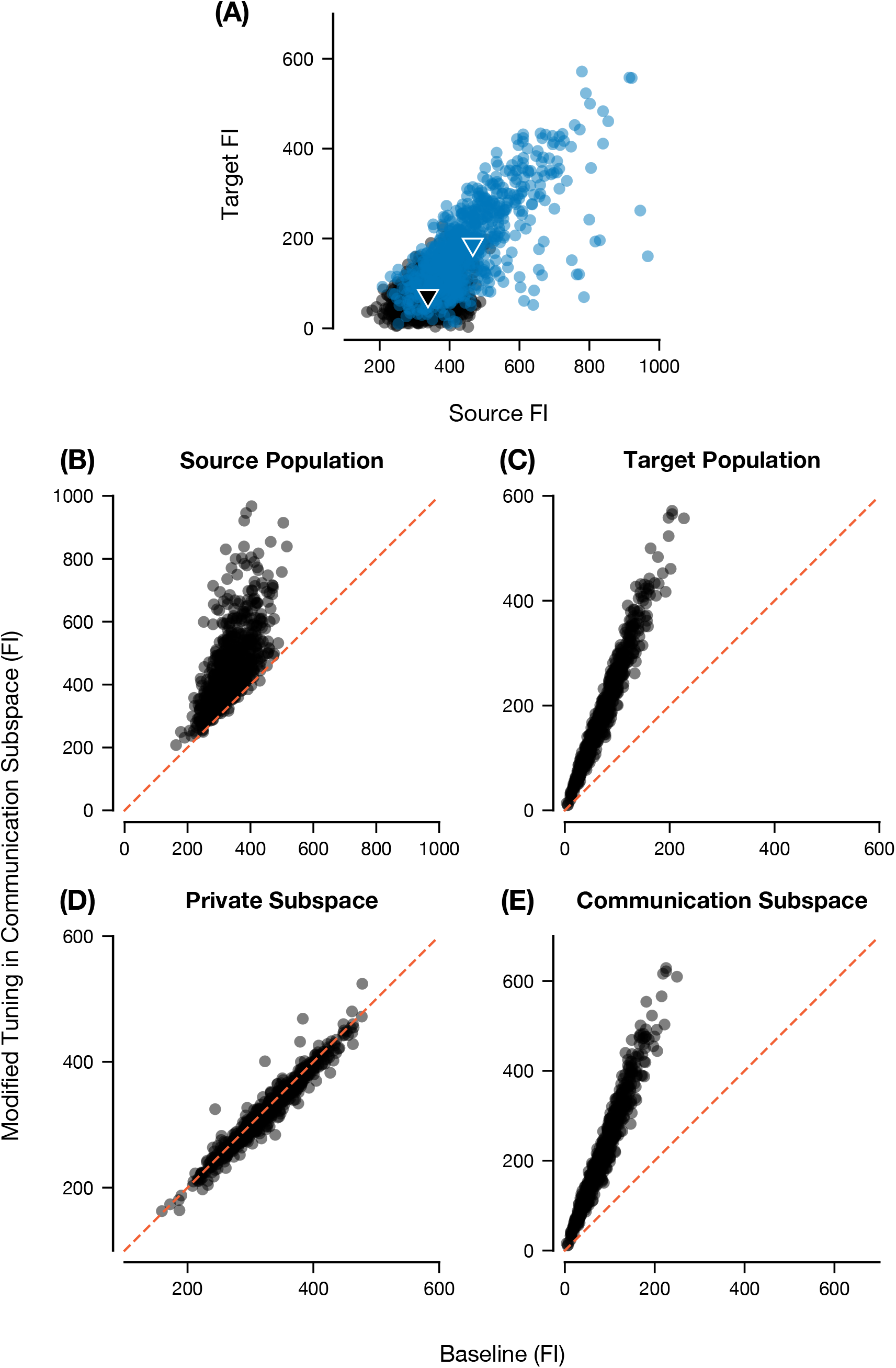
Changes in Fisher Information after Modifying Tuning and Covariance in the Communication Subspace. Same as Fig 6 except we modify the covariance matrix so that the variance is equal to the mean.

In summary, we have shown how we can selectively alter the amount of stimulus-related information communicated between the source and target areas by modifying the amount of mean stimulus tuning contributed by the private subspace (Figs 4 and 5) or communication subspace (Figs 6 and 8). We also found that the effect of these modifications on Fisher information scales with the dimensionality of the subspace where we increase tuning (Fig 7). These results demonstrate how functional networks in the brain might exploit the flexibility afforded by communication subspaces (Semedo, Zandvakili, et al. 2019) to flexibly route stimulus information between different downstream target areas.

### 3.3 Modifying Communication by Changing Target Covariance Structure

In this previous section, we investigated how to modulate communication of stimulus information by altering activity in the source area. Now, we will consider how changing activity in the target area can affect the strength of communication, which is referred to as input signal gating or selective reception (Gisiger and Boukadoum 2011; Pesaran et al. 2021; Barbosa et al. 2023). This is also related to multisensory integration, where different upstream source areas represent separate sensory modalities (e.g., vision and hearing), and a single downstream target must selectively combine information from each area (Angelaki et al. 2009). This integration of evidence requires the target area to weigh information from the upstream source areas in different ways depending on task demands (e.g., there will be scenarios when the target area will “ignore” information coming from a particular source area). Since we assumed that the target area has no residual tuning, we can only modify activity in the target area by changing the residual noise covariance or the structure of the reduced-rank regression matrix. Because changing the reduced-rank regression matrix will also change the communication and private subspaces, we will leave this fixed and focus on how changing the noise covariance in the target area can affect communication. To measure this, we will examine the relationship between the Fisher information in the communication subspace (**J**_cs_) and the impactful information in the target area (**J**(**B**_cs_**x** + ***η***_*y*_; *θ*)), which in our simulations is equivalent to the total target Fisher information (see Section 2.3). As we have previously shown, the Fisher information in the communication subspace is an upper bound for the impactful information in the target area (section 2.2.2 and AppendixE). We can interpret the tightness of this upper bound (i.e., how close **J**(**B**_cs_**x** + ***η***_*y*_; *θ*) is to **J**_cs_) as a measure of how information is being read out from the target area.

One way to weaken communication between the source and target area is to increase the noise variance in the target area (Fig 9B). In Section 2.1, we discussed how the residual variance in the target area is the quantity that is being minimized (i.e., the regression matrix is chosen such that it minimizes prediction error, which is the unexplained variance (Izenman 2008, Eq 6.78)). By increasing the variance in the target area, we are, in effect, decreasing the predictive power of the regression, such that activity patterns in the target are no longer well described as linear combinations of activity patterns in the communication subspace as they become more “random” from trial-to-trial (compare Fig 9A and Fig 9B). This decreases the amount of information decodable from the target population, explaining why the Fisher information in the target area decreases, thus “weakening” the communication between the source and target.

**Figure 9.**
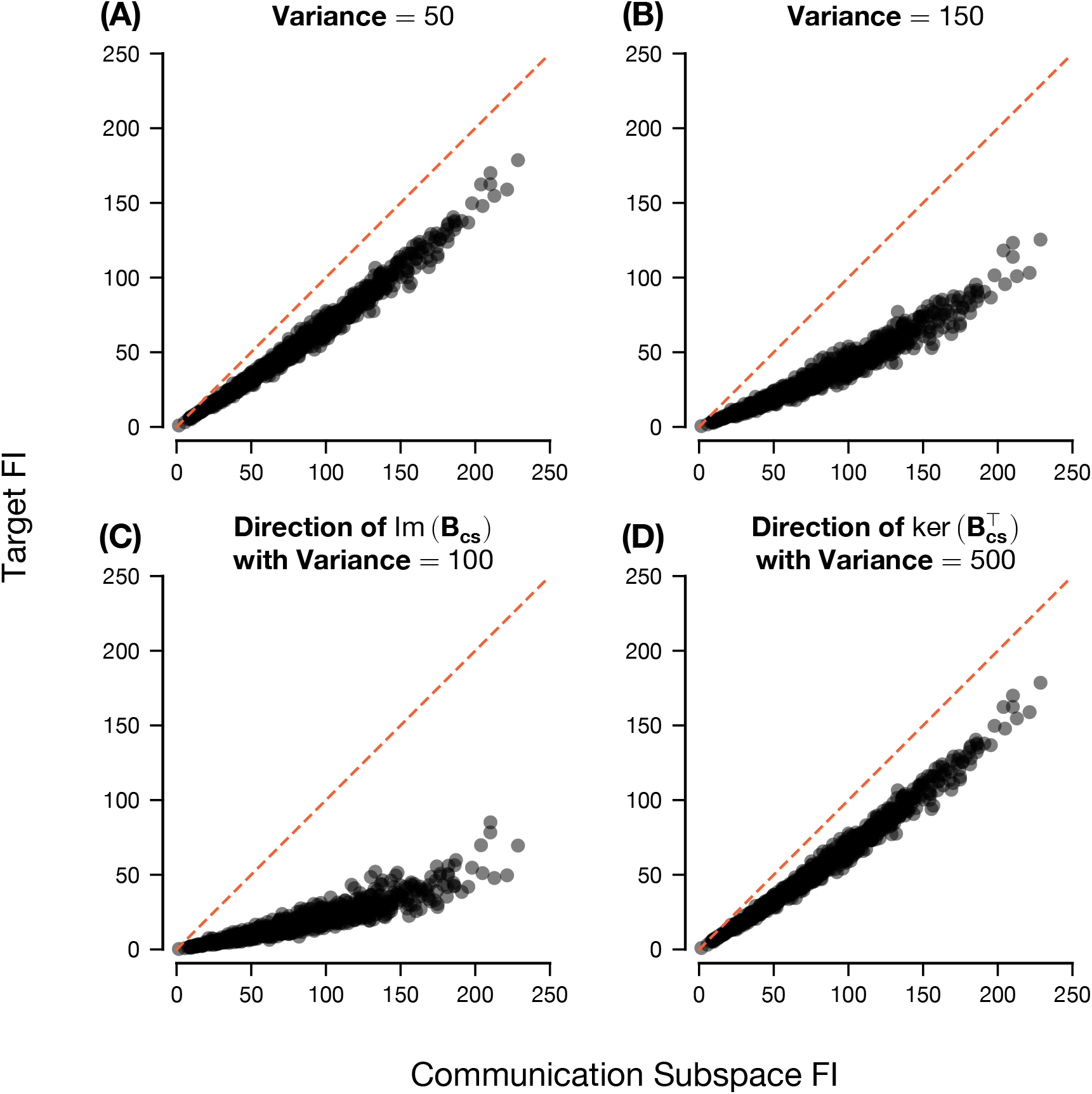
Changing Noise Covariance in Target Area can Selectively Alter Communication. Plotting the Fisher information (FI) in the communication subspace **J**_cs_ vs. the Fisher information in the target population, which for our simulations is equal to **J**(**B**_cs_**x** + ***η***_*y*_; *θ*). Note that **J**_cs_ is the same across all panels (see Eq 7). Each panel shows this relationship with a different target covariance structure 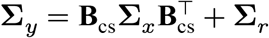 (Eq 4, with 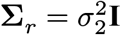). In each case, the “inherited covariance matrix” 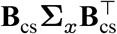 is left unchanged. Thus, each panel will have a different form of the matrix **Σ**_*r*_. (A) Baseline simulation with **Σ**_*r*_ = 50**I**. (B) Simulation with **Σ**_*r*_ = 150**I**. (C) Simulation with **Σ**_*r*_ = 50**I** + 100 Corr (Im(**B**_cs_)). To do so, we found an orthonormal basis for the column space of **B**_cs_, formed a projection matrix from this basis, converted this projection matrix to a correlation matrix and multiplied by 100. (D) Simulation with 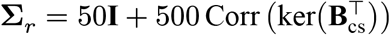. We used the same technique as in (C) but for the null space of 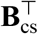.

However, this modification increases variability uniformly in every direction in the target area. In the multisensory integration scenario, in which there are two source areas, say *S*_1_ and *S*_2_, communicating via low-rank linear maps **B**_1_, **B**_2_ to one target area *T*, this would decrease information transfer from both areas. How would we modify the target area covariance to decrease communication between *S*_1_ and *T* without affecting the information transmitted from *S*_2_ to *T*? Let’s assume that **B**_1_, **B**_2_ are sufficiently low-rank such that their column spaces are linearly independent (i.e., Im(**B**_1_) ∩ Im(**B**_2_) = {0}). In this case, we can introduce noise correlations along the axes of Im(**B**_1_). Intuitively, much like increasing the variance in all directions equally (Fig 9B), increasing variability in the direction that contains the mapped activity from one source (Im(**B**_1_)) will decrease the extent to which the low-rank map determines activity in the target area (i.e., reduces the predictive power of **B**_1_). However, by selectively increasing the variability in this direction, any direction orthogonal to this subspace (which includes Im(**B**_2_)), should have no change in variability. In other words, the predictive power of the reduced-rank regression from the other source area (**B**_2_) should be left unchanged.

We can reduce this to a scenario with one source area and one target area (with corresponding map **B**_cs_) while maintaining the same core idea: if we modify the noise covariance of the target area by introducing correlations in the direction of Im(**B**_cs_) (which we call the impactful direction), we can decrease the information transfer from the source to the target, but if we modify the covariance in the orthogonal direction 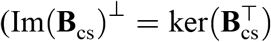, Axler 2024, Proposition 7.6), then this should not affect the information transfer. As predicted, adding noise covariance in the target area that lie along the subspace Im(**B**_cs_) decreases the impactful information in the target area relative to the Fisher information in the communication subspace (Fig 9C). Notably, this effect is more pronounced here than for the case of increasing the variability equally in all directions (Fig 9B) even though the residual covariance matrices have the same overall variance. This occurs because we are “allocating” proportionally more of the variability to the impactful directions; if we increase variability in all directions, the overall effect will depend on how this variance is projected onto the impactful directions. On the other hand, if we increase variability in the non-impactful directions (i.e., the orthogonal complement of Im(**B**_cs_)), then we see that information transmission is left unchanged, even if there is substantially stronger noise in this direction (Fig 9D).

In summary, we have shown through our Fisher information decomposition that we can modify the target activity to “silence” communication between the source and target area by introducing noise correlations in the subspace of the target area which is well predicted by the source area (Im(**B**_cs_), Fig 9C). We also showed that this effect is selective: increasing variability in any orthogonal direction does not influence how much information the target area inherits from the source area (Fig 9D).

## 4 Discussion

We have introduced a method for decomposing the stimulus-related information in cortical areas that communicate through a low-rank linear mapping (Fig 1A). We provided theoretical proofs (see Section 2.2) and numerical analyses (Figs 2 and 3) that suggest how we can interpret the individual terms (see Table 1 for descriptions of these terms) in the Fisher information decompositions. Through numerical simulations, we illustrated how to levrage communication subspaces for flexible stimulus-information routing and gating. We demonstrated how one can selectively increase the amount of information in the source area without affecting information in the target area by increasing the information in the private subspace (Figs 4 and 5). Additionally, we showed how to strengthen communication between areas by increasing information in the communication subspace (Figs 6 and 8), and that the magnitude of this effect depends on the dimensionality of the communication subspace (Fig 7). Lastly, we showed how the structure of the covariance in the target area can also influence communication by changing how much impact the information in the source area has on the target area (Fig 9).

Although we demonstrated our framework with simplified simulations using synthetic tuning and covariance, the insights gained here can guide future work on information transmission in realistic networks. For instance, we assumed no residual tuning in the target area, therefore we did not study the residual information **RI** and the synergistic/redundant information **SI**_2_. We chose not to investigate these terms to focus on several examples of how the concept of communication subspaces can be used to explain flexible information routing and signal gating. However, in experimental data, there will likely be residual tuning, which can arise from, for example, unrecorded source neurons feeding information to the target population. Future work could study these quantities by using more realistic models of activity in a source and target area (e.g., V1 and V2), for instance using hierarchical network models that reproduces the firing rate statistics of communicating neural populations (e.g., Perich and Rajan 2020; Perich, Arlt, et al. 2021; Barbosa et al. 2023; Mastrogiuseppe and Ostojic 2018; Hahn et al. 2019; Huang, Pouget, et al. 2022). Our results of Figs 4 and 6 offer concrete demonstrations of how joint changes in tuning and covariance–which are unavoidable in realistic dynamic recurrent circuit models (e.g., Seriès et al. 2004))–can impact stimulus information transmission. The framework proposed here can thus be leveraged to study quantitatively how realistic network models implement flexible or selective communication.

Despite much work on functional connectiviy, only few studies have investigated how stimulus-related information propagates between cortical areas. Panzeri and colleagues have developed a new information-theoretic measure called Feature-specific Information Transfer (FIT) to quantify stimulus-related information transfer (Celotto et al. 2023; Pica et al. 2019). Their method is based on mutual information rather than Fisher information. Although these quantities are related (Brunel and Nadal 1998; Wei and Stocker 2016), computing mutual information from neural population data is difficult due to the number of trials needed to measure the empirical multivariate probability distributions involved (Timme and Lapish 2018; Huang and Zhang 2018). As such, the FIT measure has only been applied to pairs of 1-dimensional signals (e.g. two channels in EEG recordings). In contrast, our framework has potential applicability to large population data. because, to estimate the terms in our Fisher information decompositions, one can leverage 1) techniques to efficiently compute the reduced-rank regression coefficient matrix and the associated projections (Anderson 1999; Giraud 2011; Negahban and Wainwright 2011; Hua et al. 2001); and 2) existing efficient estimators of Fisher information such as analytical bias-corrected estimation (Kanitscheider et al. 2015) and linear decoding (Kohn, Coen-Cagli, et al. 2016). As explained in Section 2.2.3, although decoding can only be applied straightforwardly to some terms in our decomposition, all the other terms can still be seen as the Fisher information for transformed measures of neural population activity (e.g., **CI**_cs_ is the linear Fisher information that corresponds to decoding the stimulus identity from **x** − **Q**_cs_**f**_*x*_).

The study most similar to ours examined the Fisher information in the output layer of a two-layer neural network (Zylberberg et al. 2017; see also Renart and van Rossum 2012). There are a few subtle differences between their work and our contribution. On the one hand, they examined how a static linear or nonlinear transformation of the output layer (i.e., the target population) changes how stimulus-related information can be decoded, while our work focuses exclusively on linear transformations. On the other hand, their analysis requires a full-rank feedforward connectivity matrix, whereas we consider low-rank connectivity. The most crucial difference, however, is the difference in perspectives that drove our analyses. Their primary concern was to understand the relationship between the capacity of a population to represent information and the ability of that population to send information to another population. Different from that work, we were not focused on what source population covariances maximize information transfer; instead, we investigated the functional implications of communication subspaces for the transmission of information (but see below for a possible extension to our framework that would consider how to optimize information transfer through a communication subspace). Nonetheless, both studies are ultimately interested in the geometry of noise correlations in neural populations and how this affects information propagation. A better understanding of this interaction through the lens of communication subspaces, possibly by combining our information decompositions with the insights of Zylberberg et al., is vital for future work.

One limitation of our proposed information decomposition is the assumption of linearity of predictability, which, in turn, leads to our analyses being based on linear geometry in Euclidean spaces. Recent work has suggested that neural population activity may lie on a “neural manifold” (Chung and Abbott 2021; Gallego et al. 2017) that can be highly nonlinear (De and Chaudhuri 2023; Fortunato et al. 2023; Golden et al. 2016; Whiteway et al. 2019). Additionally, interareal interactions may also be nonlinear (Lyamzin et al. 2015; Ruff and Cohen 2017). If these nonlinearities are significant, the partitioning of stimulus-related information we propose could significantly underestimate information transmitted between areas. Indeed, a recent simulation study demonstrated a two-layer spatial network connected via a nonlinear mapping with poor communication (measured through predictive performance of reduced-rank regression of the second layer on the first layer) even though activity in the target area is entrained to activity in the source area (Gozel and Doiron 2022). Although this is a concern, our method is still applicable even when the “true” mapping between areas is nonlinear, as long as we correctly interpret the meaning of linear Fisher information as (roughly) being the portion of the full Fisher information that is decodable through a linear estimator (see Section 2.2 for details) and if we consider a sufficiently small change in the stimulus value such that the interareal transformation is approximately linear. We can also consider possible extensions or alternatives to address nonlinear mappings between areas. The most obvious would be to combine the method proposed here with previously mentioned research on information propagation through a simple nonlinear network in which a linear weight matrix is composed with a static nonlinearity (Zylberberg et al. 2017). In such a scenario, the information in the image of such a linear-nonlinear mapping (equivalent to Eq 8) would be different than the information in the communication subspace (Eq 7), whereas for the linear mapping they are equal (Eq 30).

Ongoing research on communication subspaces indicate other possible extensions to our work. The communication subspaces framework can be seen as a kind of latent variable model (LVM) in which there are two groups of latent variables (Fig 1A): **z**_cs_ = **P**_cs_**x**, which are the latent variables that generate activity in the communication subspace, and **z**_priv_ = **Q**_cs_**x**, which are the latent variables that generate activity in the private subspace (Murphy 2021, Section 20.2). Through this interpretation, recent work has pushed the concept of communication subspaces beyond undirected, two-area communication to allow for feedforward and feedback signaling (Semedo, Jasper, et al. 2022; Gokcen, Jasper, Semedo, et al. 2022) and communication among multiple cortical areas (Gokcen, Jasper, Xu, et al. 2024). The introduction of more complicated partitioning of activity into projections onto subspaces defined by multi-area shared latent variables should be a straightforward extension of the methods we have developed here. Another kind of LVM to consider using in place of reduced-rank regression would be sufficient dimension reduction (SDR). Unlike reduced-rank regression, SDR looks for a linear map that captures the maximal amount of statistical dependency between areas (Cook 2018). By using SDR methods (e.g., Cook et al. 2015), we could find analogs of the communication and private subspaces that have minimal overlapping information, which would have some interpretational advantages over communication subspaces defined through standard reduced-rank regression (Semedo, Gokcen, et al. 2020).

An important application of our framework is the optimization of linear Fisher Information transfer. While there has been extensive work on maximizing the representation of sensory information (e.g., Brenner et al. 2000; Quian Quiroga and Panzeri 2009; Shew et al. 2011; Wang et al. 2016), research on optimally transmitting sensory information between cortical areas is in its infancy (Zylberberg et al. 2017; Ebrahimi et al. 2022; Barreiro et al. 2024; Manning et al. 2024). Our framework allows us to distinguish between communicated or received information (i.e., information in the communication subspace, Eq 7) and how this information impacts encoding in the target population (**J**(**B**_cs_**x** + ***η***_*y*_; *θ*)). Finding the communication subspace that maximizes the communicated information is essentially equivalent to finding the optimal unbiased linear decoder, implying that other axes in the source population activity space are less informative and redundant. However, this axis does not necessarily optimize how the target population “reads” the information in the communication subspace, which will depend on how the transmitted signal axis (**B**_cs_d**f**_*x*_) aligns with the target covariance matrix. This can allow for directions in the source population activity space that are not the optimal decoding axis to be more impactful on stimulus-related information in the target area. Investigating the structures of the functional maps and communication subspaces that maximize the impactful information in the target area could provide further insights into interareal communication.

## 5 Acknowledgments

We thank Adam Kohn for insightful discussion. This research was supported by NIH grants DA056400 and EY030578 (R.C.C.). This work was initiated in part at the Aspen Center for Physics, which is supported by National Science Foundation grant PHY-2210452.

## Appendix 1

To define the projections onto the communication subspace **P**_cs_ and onto the private subspace **Q**_cs_, we will use the (Moore-Penrose) pseudoinverse (written **A**^+^) (Penrose 1955). The pseudoinverse is the unique matrix satisfying the following equations:

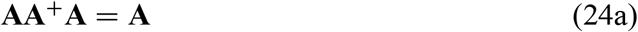

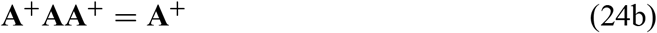

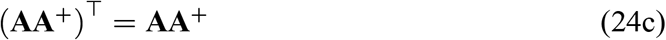

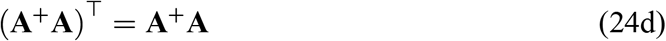

Concretely, if **A** = **USV**^⊤^ is the singular value decomposition (SVD), then **A**^+^ = **VS**^+^**U**^⊤^, where **S**^+^ is a diagonal matrix whose nonzero entries are the reciprocals of the nonzero diagonal entries of **S** (i.e., the reciprocals of the nonzero singular values of **A**; the singular values equal to 0 are retained). For any linear map **A**, the map **A**^+^**A** is equivalent to the projection onto ker(**A**)^⊥^ (Axler 2024, Propositions 6.69 and 7.75). As this is how we defined the communication subspace previously, we can define the projection onto the communication subspace as

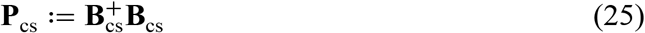

## B Appendix 2

Here we prove that the two expressions for Fisher information in the communication subspace, introduced in Section 2.2.1), are equivalent, using the following proposition:

### Lemma 1.

*Let* **A, B** *be injective matrices (i*.*e*., *matrices with full column rank) of size n* × *k, and let* **C** *be a k* × *k square invertible matrix. For any k-dimensional vectors a, b, the following identity holds:*

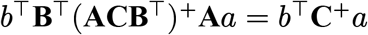

*Proof*. It suffices to show that (**ACB**^⊤^)^+^= (**B** ^⊤^)^+^**C**^+^**A**^+^ since

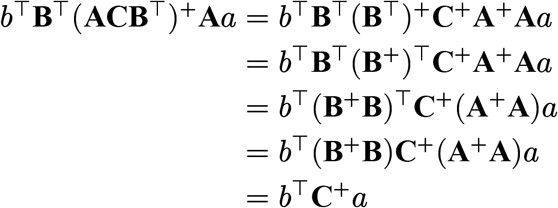

Where we have used the facts that **B**^+^**B** is symmetric (Eq 24d) and the pseudoinverse of an injective matrix is a left inverse of **A** (i.e., **A**^+^**A** = **I** for **A** injective) because 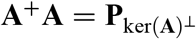 and ker(**A**) = {0} since **A** is injective; thus, its orthogonal complement is the whole space. In order to show that (**ACB**^⊤^)^+^ = (**B**^⊤^)^+^**C**^+^**A**^+^ we simply check that (**B**^⊤^)^+^**C**^+^**A**^+^ satisfies Eqs 24a-d for **ACB**^⊤^; by uniqueness of the pseudoinverse, the result follows. To see this, we first note that:

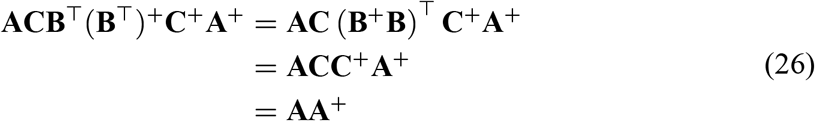

Because **B** is injective and **C** is invertible. This proves that (**B**^⊤^)^+^**C**^+^**A**^+^ satisfies Eq 24c, because **AA**^+^ is symmetric by the same requirement. A similar computation shows that (**B**^⊤^)^+^**C**^+^**A**^+^**ACB**^⊤^ = **BB**^+^, which shows that (**B**^⊤^)^+^**C**^+^**A**^+^ satisfies Eq 24d because **BB**^+^ is symmetric (Eq 24c). Using these facts, we can calculate:

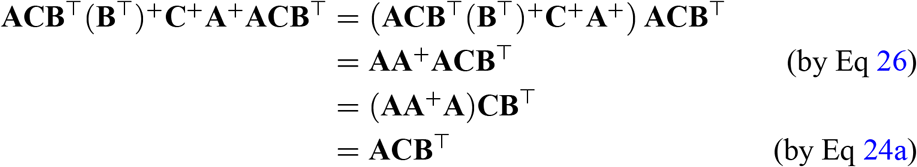

Which shows that (**B** ^⊤^)^+^**C**^+^**A**^+^ satisfies Eq 24a. Similarly, to see that this meets Eq 24b,

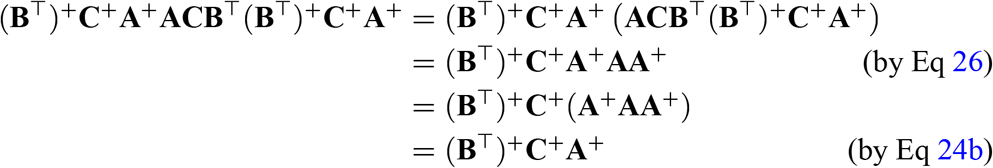

Therefore, (**B**^⊤^)^+^**C**^+^**A**^+^ meets all of the requirements to be a pseudoinverse of **ACB**^⊤^, so (**ACB**^⊤^)^+^ = (**B**^⊤^)^+^**C**^+^**A**^+^.

We additionally need the following three facts: first, **Σ** is symmetric positive semidefinite (that is, if *x*^⊤^**Σ***x* ≥ 0 for all vectors *x*) *if and only if*

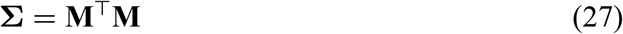

If **Σ** is also invertible (i.e., symmetric positive-*definite, x*^⊤^**Σ***x* > 0 for all vectors *x* ≠ 0), then **M** in Eq 27 can be taken to be a symmetric and invertible matrix (Zhang 2011, Theorem 7.3).

Second, for any matrix **C**,

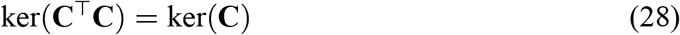

To see this, suppose *x* ∈ ker(**C**^⊤^**C**). Consider *x*^⊤^**C**^⊤^**C***x* = *x*^⊤^0 = 0 = (**C***x*)^⊤^(**C***x*). By definition of the inner product, this implies **C***x* = 0, or *x* ∈ ker(**C**), so ker(**C**^⊤^**C**) ⊆ ker(**C**), and the other inclusion is trivial.

Finally, if **Σ** is a positive-definite matrix and **V** is an injective matrix, then **A** = **V**^⊤^**ΣV** is positive-definite. To see this, let **Σ** = **M**^2^ by Eq 27. Then

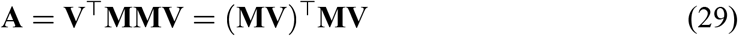

so **A** satisfies Eq 27, which means that **A** is positive semi-definite. To show that **A** is positive-definite, we need to show that **A** is invertible. To do so, it suffices to show that ker(**A**) = {0}, since **A** is a square matrix and by the rank-nullity theorem, ker(**A**) = {0} implies **A** has full column-rank. Suppose *x* ∈ ker(**A**) = ker(**V**^⊤^**ΣV**). This implies that *x* ∈ ker(**MV**) because **A** is positive semi-definite (Eq 29) and by Eq 28. In other words, this means **MV***x* = 0. Because **M** is invertible, this implies that **V***x* = 0. However, this means *x* = 0 by the injectivity of **V**, so ker(**A**) = {0}.

Now, we can prove that

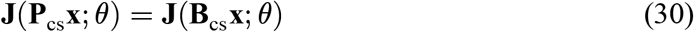

*Proof*. For **B**_cs_, we define the compact SVD by 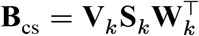, for **S**_*k*_ a diagonal *k* × *k* matrix corresponding to the non-zero singular values of **B**_cs_. In this notation, 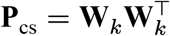 by direct computation.

We will first show that 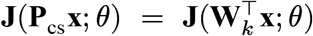 Define 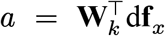 and 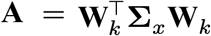 From the above, **A** is positive-definite and, in particular, invertible. Then

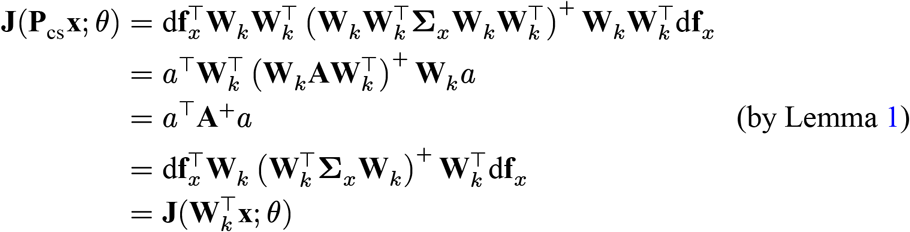

Now, it suffices to show that

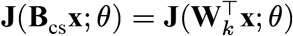

We can do so using the same logic as above. Define *a*, **A** as before, and let 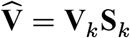. Note that 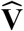 is an injective matrix: if **V**_*k*_**S**_*k*_*x* = 0, by the injectivity of **V**_*k*_ this imples **S**_*k*_*x* = 0. However, **S**_*k*_ is invertible, so this is only true if *x* = 0. Then, using the SVD of **B**_cs_:

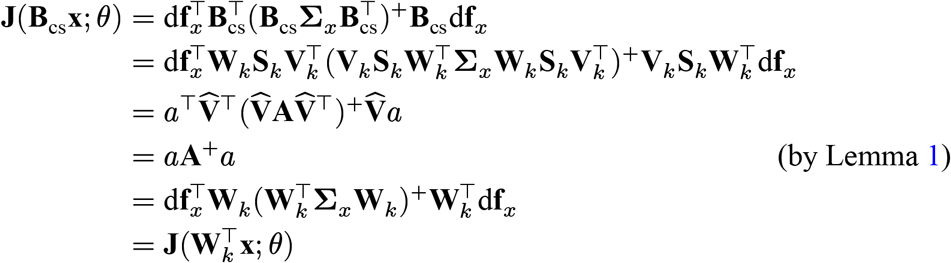

## C Appendix 3

### Lemma 2.

*Suppose* **Σ** *is a symmetric positive-definite matrix and let* **P** *be a orthogonal projection (***P**^2^ = **P** *and* **P**^⊤^ = **P***). Then*

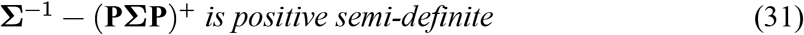

*Proof*. Let *n* be the rank of **Σ** and let *k* < *n* be the rank of **P**. There exists a basis for ℝ^*n*^ such that 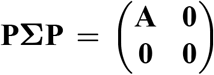 for **A** a symmetric positive-definite *k* × *k* matrix: create an orthonormal basis {*u*_1_, …, *u*_*k*_} for the column-space of **P** and complete to a basis **U** of ℝ^*n*^; the matrix **U**^−1^**PU** is now of the form 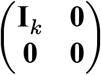, so **PΣP** has the desired form in this basis.

Similarly, in this basis 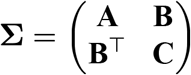, for **B** a *k* × *n* − *k* matrix and **C** an *n* − *k* × *n* – *k* matrix. We can write **Σ**^−1^ as a function of the matrices in this partition using the Schur complement **Σ**/**A** = **C** − **B**^⊤^**A**^−1^**B** (Zhang 2011, Theorems 2.3 and 2.4) and compute 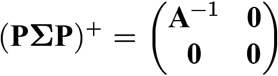 directly by checking the defining properties of the pseudoinverse (Eq 24). With this notation:

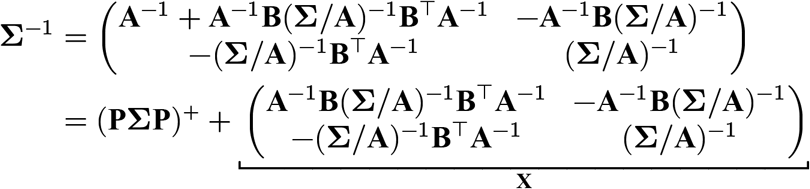

It suffices to show that **X** defined above is positive semi-definite. To show this, we need to find a matrix **Y** such that **X** = **Y**^⊤^**Y** (Eq 27). Because **Σ** is positive-definite, **Σ**/**A** is also positive-definite (Gallier 2011, Proposition 16.2), which implies (**Σ**/**A**)^−1^ is positive-definite and there is a symmetric matrix **M** such that (**Σ**/**A**)^−1^ = **M**^2^. Define **Y** = (−**MB**^⊤^**A**^−1^ **M**), which is an *n* − *k* × *n* matrix. Then

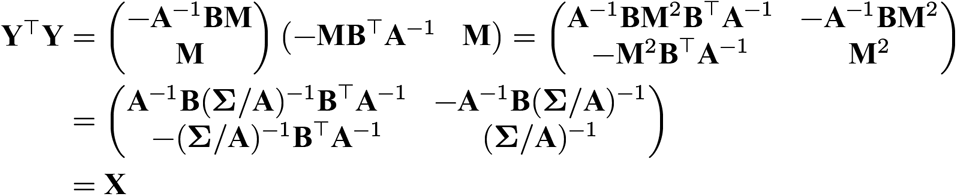

Therefore, **X** = **Σ**^−1^ − (**PΣP**)^+^ is positive semi-definite.

To see how this proves Eq 16, write:

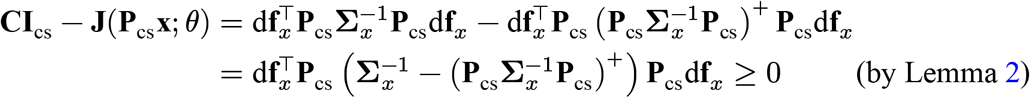

and similarly for Eq 17. As noted, this also shows that **J**(**x**; *θ*) ≥ **J**_cs_ and **J**_priv_, as similar logic to the above proof would demonstrate that **Σ**^−1^ − **P** (**PΣP**)^+^ **P** is positive semi-definite.

## D Appendix 4

Let υ = **P**_cs_d**f**_*x*_ and *w* = **Q**_cs_d**f**_*x*_. Then

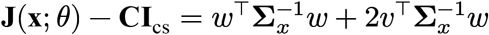

There must be some υ such that 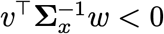 (for example, −*w* is one such vector because 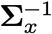 is positive-definite, so 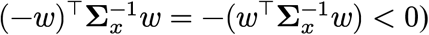 Let 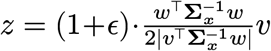 for *ϵ* > 0 some constant. Then since **Σ**_*x*_ is positive-definite, we have

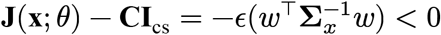

This calculation shows that there will be some tuning curve and some population covariance matrix such that **J**(**x**; *θ*) < **CI**_cs_.

## E Appendix 5

The more direct argument to prove that the impactful information is bounded by the information in the communication subspace, would be to show, similar to Lemma 2, that

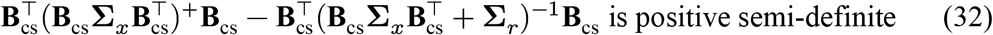

To do so, we prove the following lemma (this lemma and the argument that this proves Eq 32 are due to user1551 n.d.).

### Lemma 3.

*Let* **A** *be an n* × *n symmetric positive semi-definite matrix, let* **B** *be an n* × *n symmetric positive-definite matrix, and assume that* **A** + **B** *is invertible. Then*

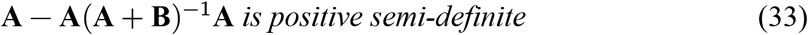

*Proof*. By assumption, **A** is positive semi-definite, so it trivially follows that **A** + **B** − **B** is positive semi-definite. Because **A** + **B** and **B** are positive-definite and invertible, we have that **B**^−1^ −(**A** + **B**)^−1^ is positive semi-definite (Zhang 2011, Theorem 7.8). For any positive semi-definite matrix **X** and any matrix **Y, Y**^⊤^**XY** is positive semi-definite (Eq 29), and **B** is symmetric. This means that

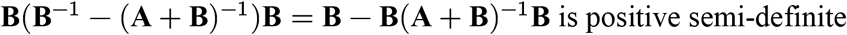

By defining **C** = **A** + **B**, we can rewrite this in terms of **A** and **C**:

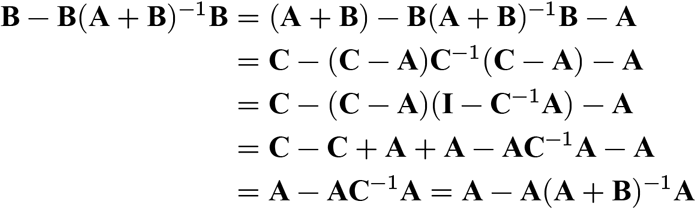

Therefore, **A** − **A**(**A** + **B**)^−1^**A** positive semi-definite.

To see how this generalizes Eq 32, we prove that 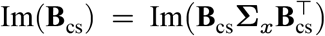. The inclusion 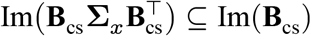 is trivial. To prove that they are equivalent, we show that the images have the same dimensionality, which implies that they are equal because the only *n* dimensional subspace of an *n* dimensional vector space is itself. For this, we consider 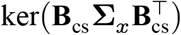. Because **Σ**_*x*_ is symmetric positive-definite, **Σ**_*x*_ = **M**^2^ for **M** symmetric and invertible (Eq 27). Using this and Eq 28, we find that

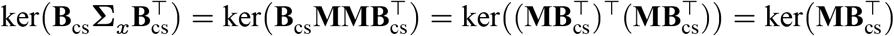

Since **M** is invertible, 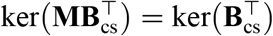, so we have 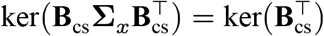.

By using the rank-nullity theorem and the fact that row-rank is equivalent to column-rank 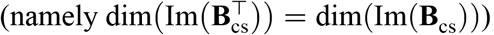

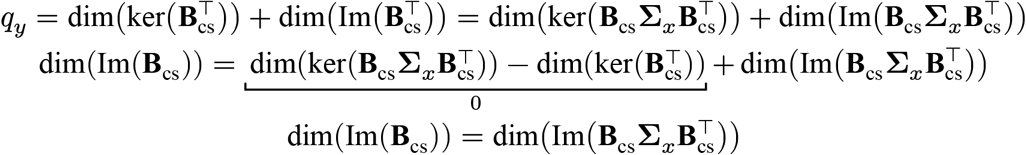

Therefore, because Im(**B**_cs_) and 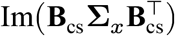 have the same dimensionality and 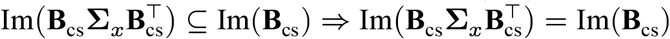.

Now, since **B**_cs_d**f**_*x*_ ∈ Im(**B**_cs_) trivially, there is some vector υ in the target population activity space such that 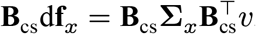 Then

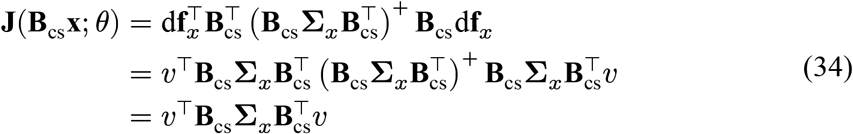

Where the last line of Eq 34 follows from a defining property of the pseudoinverse (Eq 24a).

Similarly, we expand **J**(**B**_cs_**x** + ***η***_*y*_; *θ*) as

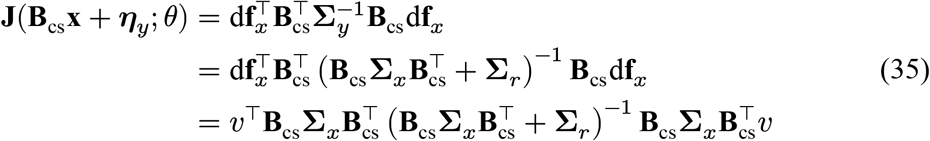

Using Eqs 34 and 35, we can restate Eq 19 as

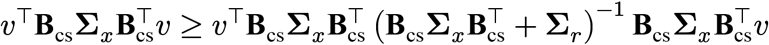

After rearranging this, we have

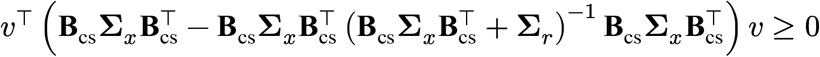

In other words, Eq 19 is equivalent to the claim that

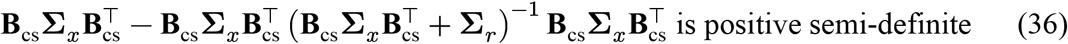

Now defining 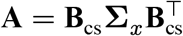 and **B** = **Σ**_*r*_ in Eq 36, we get Eq 33. Therefore, as a corollary of Lemma 3, we have proved Eq 32 and therefore Eq 19, namely that the information in the communication subspace **J**_cs_ = **J**(**B**_cs_**x**; *θ*) is an upper bound on the impactful information **J**(**B**_cs_**x** + ***η***_*y*_; *θ*).

## F Appendix 6

Here we derive mathematically the results illustrated by the simulations of Fig 4. Central to these derivations will be the following identities:

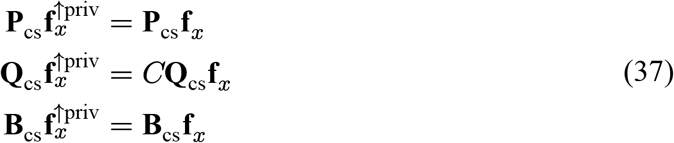

These follow immediately from the basic properties of orthogonal projections applied to Eq 22. First, we will use the source Fisher information decomposition (Eq 12) for **J**(**x**^↑priv^; *θ*) and simplify using these identities to obtain:

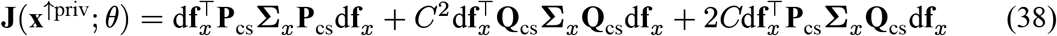

If we examine the difference between the source Fisher information before and after modifying the private tuning

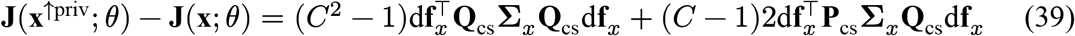

Defining 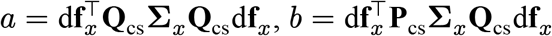, simple algebra will show that this difference is positive as long as 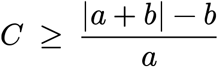. This shows that the source Fisher information will increase after modifying the private tuning in this manner (Fig 4B).

Next, if we look at the target Fisher information decomposition (Eq 13 with **r**_*y*_ = 0) and using the identities (Eq 37), we obtain

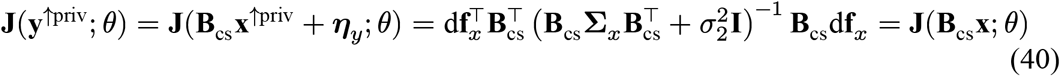

This shows that there is no change in the amount of Fisher information in the target area. The same derivation applies to the information in the communication subspace (using Eq 40 with *σ*_2_ = 0 and using the pseudoinverse), also demonstrating that the Fisher information in the communication subspace will remain invariant to changes of tuning in the private subspace (Fig 4E).

Lastly, if we compare the Fisher information in the private subspace for **x** and **x**^↑priv^, **J**(**Q**_cs_**x**^↑priv^; *θ*) − **J**(**Q**_cs_**x**; *θ*), we can write this difference as

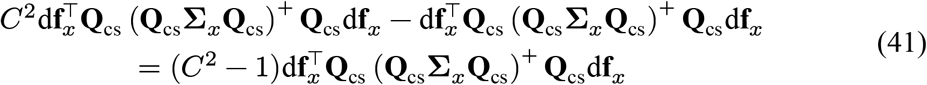

Thus, as long as *C* > 1, then **J**(**Q**_cs_**x**^↑priv^; *θ*) ≥ **J**(**Q**_cs_**x**; *θ*), or the Fisher information in the private subspace will increase (Fig 4D).

## Footnotes

1 This paper will use the words “area” and “population” interchangeably. When one considers *intra-areal* communication between subpopulations in the same region of cortex using the same techniques used to characterize interareal communication, it is more correct to use “population” (e.g., the “source population”).

2 Strictly speaking, the information in the private subspace is not a component of the source Fisher information; instead, the *contributed information* to the private subspace (Eq 15) is a component of the source Fisher information and the information in the private subspace is merely a lower bound for both the source Fisher information (Eq 11) and the contributed information (Eq 17). Nonetheless, we provide a mathematical justification for the fact that increasing the information in the private subspace tends to increase the information in the source area (Eq 39).

3 If −*b* > *a*, then 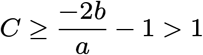 otherwise, *C* ≥ 1. Note that we ignore the smaller root of Eq 39, but in general, if we choose *C* to be less than this smaller root, then Eq 39 will also be positive.

